# *H_2_O*: Improving Detection in Large-Scale Gene Family Expansions and Associated Phylogenomic Inferences

**DOI:** 10.64898/2026.11.20.768191

**Authors:** Keyi Feng, Michelle L. Gaynor, Stephen A. Smith

**Author notes:** Corresponding Author: Keyi Feng.

## Abstract

Ancient whole-genome duplications (WGDs) have shaped flowering plant evolution, but their placement is sometimes uncertain, especially in clades primarily studied through transcriptomes. In such datasets, peaks in synonymous substitutions among paralogs (Ks) and phylogenomic gene-duplication mapping can identify large-scale gene family expansions (GFEs), but these approaches may disagree and, on their own, cannot distinguish WGD from chromosomal duplication or other genome-scale duplication mechanisms. These discrepancies may stem from biological processes such as diploidization, hybridization, and incomplete lineage sorting, or from systematic biases introduced by hidden gene paralogy that current pipelines fail to address. Here, we present *H_2_O* (homolog-to-ortholog), a workflow that integrates Ks distributions with tree-based conservative gene duplication calls (node- partitioned Ks distributions) and orthology inference with duplication-informed data-filtering methods. It improves the detection and placement of transcriptome-inferred GFEs while reducing errors from hidden paralogy. We evaluated *H_2_O* using simulated datasets and three empirical datasets (Ericales and two Caryophyllales subclades), refining large-scale GFE placements and comparing phylogenomic outcomes with those of ASTRAL-Pro and Yang and Smith (2014). Across empirical datasets, node-partitioned Ks distributions improved confidence in large-scale GFE placement and, under conservative phylogenetic data filtering, significantly reduced associated gene tree discordance. Reduced discordance comes at the cost of substantial data removal. Through simulation, we demonstrated that *H_2_O*’s data-filtering methods effectively removed all paralogous relationships that orthology inference methods sometimes report, reducing gene tree discordance. These results highlight the need to treat transcriptome-inferred GFEs cautiously, integrate multiple lines of evidence, and explicitly account for hidden paralogy in plant phylogenomics.

## Introduction

Large-scale gene family expansions (GFEs), including ancient whole-genome duplications (WGDs), are among the most influential evolutionary processes in the history of flowering plants (Soltis et al. 2014; Landis et al. 2018; Heslop-Harrison et al. 2023). Recent studies have found putative WGDs to be pervasive across angiosperm lineages (Ren et al. 2018; Leebens- Mack et al. 2019) and have been linked to numerous major trait innovations and ecological shifts, including the origin of the flower, the diversification of core eudicot floral structures, the evolution of carnivory in Caryophyllales, climatic niche shifts, and improved stress tolerance (Chanderbali et al. 2016; Walker et al. 2017; Clark and Donoghue 2018; Smith et al. 2018b; Sobhanian et al. 2026).

These genomic duplications are identified and analyzed based on assembled genomes and transcriptomes. High-quality, fully assembled genomes enable effective WGD identification deep in angiosperm evolutionary history through synteny and phylogenomic analyses, despite WGD signal degradation by diploidization (Tang et al. 2008; Amborella Genome Project 2013; Jiao et al. 2014; Feng et al. 2026). However, as reference genomes are limited at the order level and below, transcriptomes serve as an alternative. Transcriptomes are relatively easy and inexpensive to generate; thus, they serve as the primary data source for detecting GFEs at lower taxonomic levels (Huang et al. 2016; McKain et al. 2016; Cai et al. 2019; Yang et al. 2018; Carruthers et al. 2024). Yet, despite extensive research, substantial uncertainty remains in accurately identifying GFEs using transcriptomic data. Without precise identification and phylogenetic placement of large-scale GFEs, it remains challenging to further explore the evolutionary consequences of these important events.

Large-scale GFEs detected from transcriptomes are often treated as putative WGDs. Because synteny analyses are not feasible with transcriptomes, the distribution of synonymous substitutions between paralog gene pairs (Ks) has become one of the most common methods for detecting putative WGDs in transcriptomic studies, usually in conjunction with phylogenomic inferences and chromosome counts (Cai et al. 2019; Ren et al. 2018; Yang et al. 2018; Zhao et al. 2023; Feng et al. 2024; Carruthers et al. 2024). Ks values reflect the amount of molecular evolution between paralogous gene pairs and have been used as a proxy for the time since gene duplication. Under this framework, a peak in the Ks distribution indicates evidence of a putative WGD, a simultaneous burst of many paralogous genes at a single time point.

Similarly, in phylogenomic inference, gene duplications are identified by mapping thousands of homologous phylogenetic trees reconstructed from transcriptomes onto a consensus species tree (Smith et al. 2015; McKain et al. 2016; Zwaenepoel and Van de Peer 2019; Chen et al. 2022). Gene duplications that cluster at specific nodes of the consensus species tree are used to suggest putative WGDs. Further support for WGD can be seen in base chromosome number shifts, which may indicate changes in genome size or subsequent genome rearrangements.

While transcriptomes are one of the most popular data sources for studying angiosperm genomic evolution, they alone are insufficient to confirm WGDs because they lack syntenic information. For example, genomic syntenic analyses have shifted the traditional view of the polyploid events that the model polyploid grass, *Sporobolus alterniflorus*, has undergone (Salmon et al. 2025). Previously, *S. alterniflorus* was believed to have experienced three ancestral WGDs, but genomic synteny analysis revealed that it is a diploidized meso-octoploid or neotetraploid that experienced two ancestral WGDs and a base chromosome number shift. There is no definitive evidence of ancestral WGD without synteny analyses of assembled genomes in conjunction with an increase in base chromosome number. What Ks distributions and phylogenomic gene-duplication inferences capture are the signals of large-scale GFEs.

Without syntenic information, these two methods cannot differentiate the sources of these large-scale GFEs from WGD, chromosomal duplication, segmental duplication, or other mechanisms. Therefore, we define bursts of gene duplications supported by Ks distributions and phylogenomic analyses, but lacking other evidence, as large-scale GFEs. Large-scale GFEs with a near-doubling of the base chromosome count in the descendant clade, and confirmed by other synteny studies, are considered putative WGD events. It is essential to determine what the available methods can detect before exploring the methodologies.

Although methods for detecting large-scale GFEs using transcriptomes are well established, there are two major sources of uncertainty in their detection. First, uncertainty in large-scale GFE estimation arises from treating Ks distributions and phylogenomic gene-duplication inferences as separate lines of evidence. Existing WGD detection methods either utilize only one of the two (McKain et al. 2016; Zwaenepoel and Van de Peer 2019; Chen et al. 2024), or analyze them separately (Yang et al. 2018; Chen et al. 2022). Although both methods identify the same phenomenon, their results do not consistently align owing to unidentified factors (Stull et al. 2021; Yang et al. 2018; Carruthers et al. 2024; Feng et al. 2024). The mean of Ks peaks varies around a given event, and the presence or absence of a peak does not constitute definitive evidence of a GFE/WGD (Tiley et al. 2018). Ks peaks represent simultaneous bursts of gene family expansion within each species, but whether peaks with similar means across different taxa represent the same GFE event remains uncertain. In contrast to Ks distributions, phylogenomically inferred gene duplications that cluster at a node in the consensus species tree suggest that this group of species shares many GFEs, but whether these GFEs originated simultaneously is unknown. Only when many gene-family expansions are inferred to have occurred near the same evolutionary time interval and are shared by a lineage do we interpret the signal as a large-scale GFE. Ideally, for a putative large-scale GFE event, Ks peaks are shared by the same groups of species as those with many gene duplications mapped to the same node phylogenetically (Figure 1a). In practice, Ks peaks and bursts of gene duplication sometimes correspond to neighboring but different nodes, obscuring the placement of the event (Figure 1b). Because Ks distributions and phylogenomic gene-duplication inferences provide complementary information about GFEs, combining these two methods in a single analysis should, in principle, reconcile some discrepancies (Figure 1c). This consideration is currently lacking in existing WGD detection methods.

**Figure 1.**
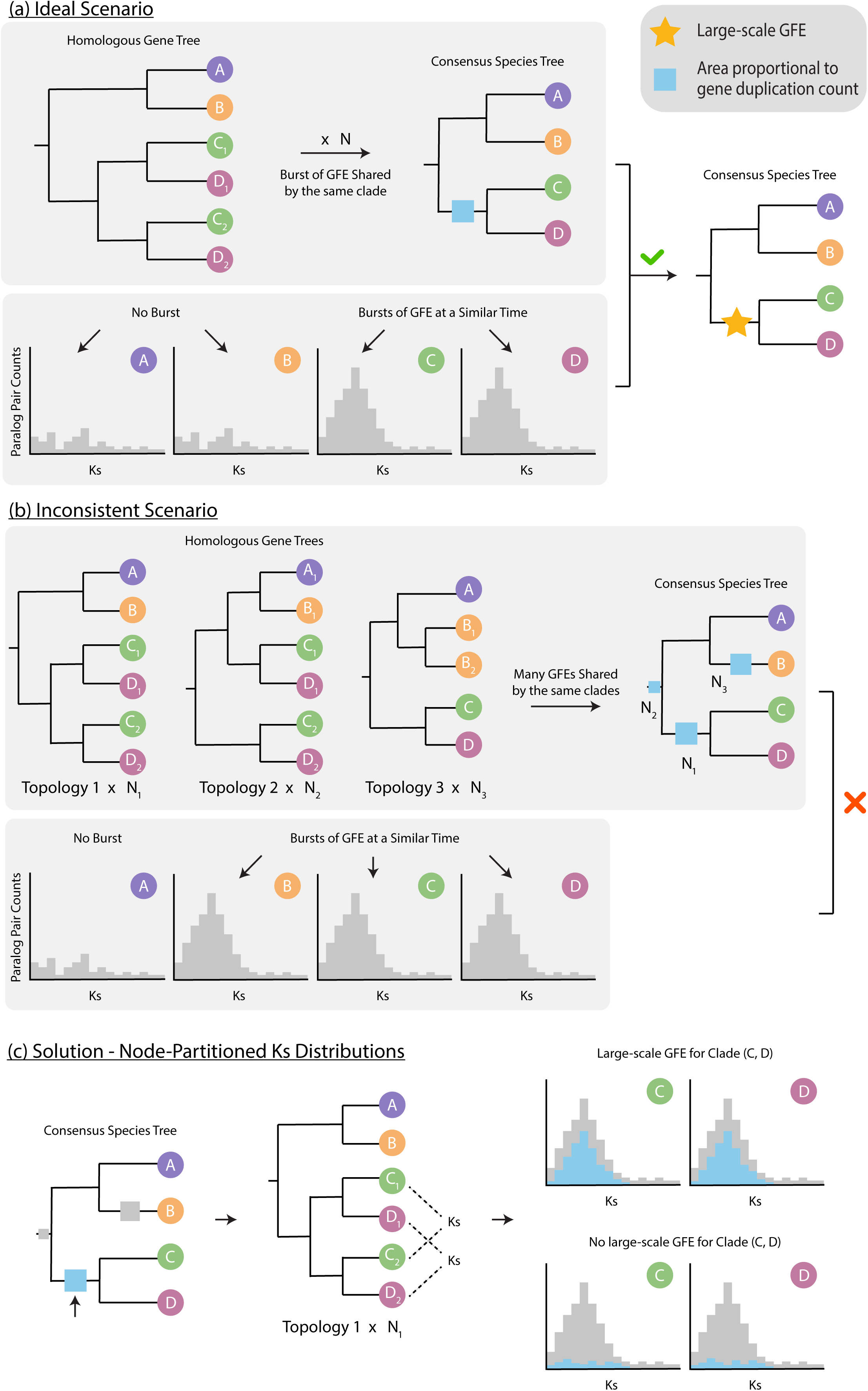
Large-scale gene family expansion (GFE) identification based on the consensus of phylogenomic inference of gene duplications and Ks distributions. In the phylogenies, colored circles represent gene sequences for species A-E; subscript numbers indicate duplicated gene copies. “N” refers to a certain number. (a) An ideal scenario for clade ((A, B), (C, D)). The two methods are separated into two gray boxes; gene-duplication inference is on top. Species C and D have Ks peaks with a similar mean, while A and B do not have peaks in the distribution. A large number of gene duplications inferred from homolog trees are mapped to the MRCA of C and D. Both lines of evidence agree and support the placement of a large-scale GFE before the speciation event that separated C and D. (b) An inconsistent scenario that is simplified from empirical data for the same hypothetical clade. Only B, C, and D have Ks peaks with similar means, while many gene duplications are inferred from neighboring nodes. The Ks and phylogenomic gene-duplication mapping results do not align, making it difficult to determine the underlying evolutionary event. (c) Proposed solution to connect Ks and phylogenomic inference with node-partitioned Ks distributions. For each node with a large number of mapped gene duplications, calculate Ks for all the paralog gene pairs duplicated at that node, and highlight those Ks values (blue) in the Ks distribution (gray is all Ks values for that species). Alt text: The figure has three parts, a, b, and c, each presenting a scenario for identifying large- scale gene family expansion (GFE) using phylogenomic gene-duplication inference and Ks distributions for four species: A, B, C, D. In panel a, for gene duplication inference, this panel first illustrates N numbers of homolog gene trees of ((A, B), ((C_1_, D_1_), (C_2_, D_2_))), corresponding to N gene duplications mapped before the most recent common ancestor (MRCA) of species C and D in the consensus species tree. This matches the Ks distributions described in the caption. In panel b, for gene duplication inference, it displays N_1_ homolog gene trees of ((A, B), ((C_1_, D_1_), (C_2_, D_2_))), N_2_ trees of (((A_1_, B_1_), (C_1_, D_1_)), (C_2_, D_2_)), and N_3_ trees of ((A, (B_1_, B_2_)), (C, D))), corresponding to N_1_ gene duplications mapped before (C, D), N_2_ duplications before ((A, B), (C, D)), and N_3_ duplications before species B in the consensus species tree. This does not match the Ks distributions described in the caption of panel b. In panel c, we illustrate a solution to the discrepancy in panel b: focusing on each specific node (C, D), in this case, Ks values for all paralog pairs duplicated at this node are highlighted (in blue) within the gray Ks distribution. The gray distribution represents the full Ks distribution for each species. When the blue histogram forms a peak, it supports a large-scale GFE for the clade (C, D); if the blue histogram is mostly close to zero, it does not support a large-scale GFE.

Second, systematic errors in phylogenetic inferences of gene duplication and gene orthology can also lower confidence in large-scale GFE placement. Although the concept of orthology has been proposed since 1970 (Fitch 1970), its consequences for large-scale GFE detection remain underexplored. Many large-scale GFE events result from ancient hybridization, i.e., allopolyploidy (Wang et al. 2019a; Stull et al. 2023; Gaynor et al. unpublished data). Consistent with this, large-scale GFEs often coincide with elevated gene tree discordance in neighboring nested nodes in the consensus species tree, commonly attributed to biological processes such as incomplete lineage sorting (ILS) and hybridization (Triplett et al. 2014; García et al. 2017; Yang et al. 2018; Feng et al. 2024). The combination of large-scale GFE, genomic restructuring/diploidization, and ILS/hybridization can result in systematic errors in gene duplication and orthology determination due to hidden gene paralogy (Galtier and Daubin 2008; Degnan and Rosenberg 2009; Huang et al. 2016). When ILS/hybridization occurs concurrently with large-scale GFEs, gene duplication location and gene orthology may be misidentified; in cases where a duplicated gene family is reduced to a single-copy state, the gene copies may be hidden paralogs (both scenarios are detailed in the methods section).

However, systematic error and hidden paralogy have received limited attention in explaining the connection between large-scale GFEs and gene tree discordance.

ASTRAL-Pro (Zhang et al. 2020; Tabatabaee et al. 2023; Zhang et al. 2025) and Yang and Smith (2014) are two major tree-based orthology inference methods favored in plant phylogenomic research. To infer orthology from homologous gene trees, these methods need to identify duplication nodes on the homologous gene tree, similar to tree-based gene duplication mapping methods. After calling gene duplications, Yang and Smith (2014) dissect the homologous gene tree into orthologous gene trees and use them to estimate a consensus species tree, whereas ASTRAL-Pro (Zhang et al. 2025) directly uses labeled homologous gene trees to derive the consensus species tree. Although both are effective tools for tree-based orthology inference, they do not address hidden paralogy caused by the combination of large- scale GFE and ILS.

Here, we explore ways to improve the detection and placement of large-scale GFEs and minimize systematic errors in phylogenomic inferences. We present homolog-to-ortholog (*H_2_O*), a tree-based orthology inference method designed to address challenges in plant phylogenomic datasets affected by large-scale GFEs. *H_2_O* incorporates algorithms that directly incorporate phylogenetically inferred gene duplications into Ks analyses to improve large-scale GFE detection (node-partitioned Ks distributions; Figure 1c), and it employs phylogenetic data- filtering methods to address major sources of hidden gene paralogy and reduce systematic errors in phylogenomic orthology inference. We assess *H_2_O* using homolog tree datasets simulated with SimPhy (Mallo et al. 2016) and published phylotranscriptomic datasets for Ericales (Carruthers et al. 2024) and two Caryophyllales subclades (Yang et al. 2018; Feng et al. 2024), with known uncertain large-scale GFE placement and linked-gene-tree discordance.

Additionally, we compare *H_2_O’s* orthology inference outcome with ASTRAL-Pro3 (Zhang et al. 2025) and the Yang and Smith (2014) pipeline. We aim to (1) increase confidence in the detection and placement of large-scale GFEs, (2) reduce systematic errors and hidden paralogy linked to gene duplication and orthology inference in phylogenomics through data filtering, and (3) gain new insights into the genomic impacts of large-scale GFEs. Accurate inference of large- scale GFEs and associated phylogenetic relationships provides a more reliable evolutionary framework for downstream analyses, enabling more robust investigations of trait evolution, gene family expansion, and other genomic consequences of gene duplication.

## Materials and Methods

### H_2_O’s Workflow

*H_2_O*’s two main functionalities are phylogenetic ortholog inference and gene-duplication mapping based on input homolog trees (Figure 2a; https://github.com/fkyeeb/H2O). For ortholog inference, it implements a standard approach and two special filtering methods to reduce hidden paralogy. The workflow can be divided into three steps: (1) the standard orthology inference, which must be completed before we can proceed to gene-duplication mapping and the special filtering steps; (2) gene-duplication mapping via *H_2_O* and large-scale GFE identification with external analyses or information; (3) homolog and ortholog special filtering methods, “pruning” and “GFE-filtering”, to reduce hidden paralogy and systematic errors in orthology inference related to large-scale GFEs.

**Figure 2.**
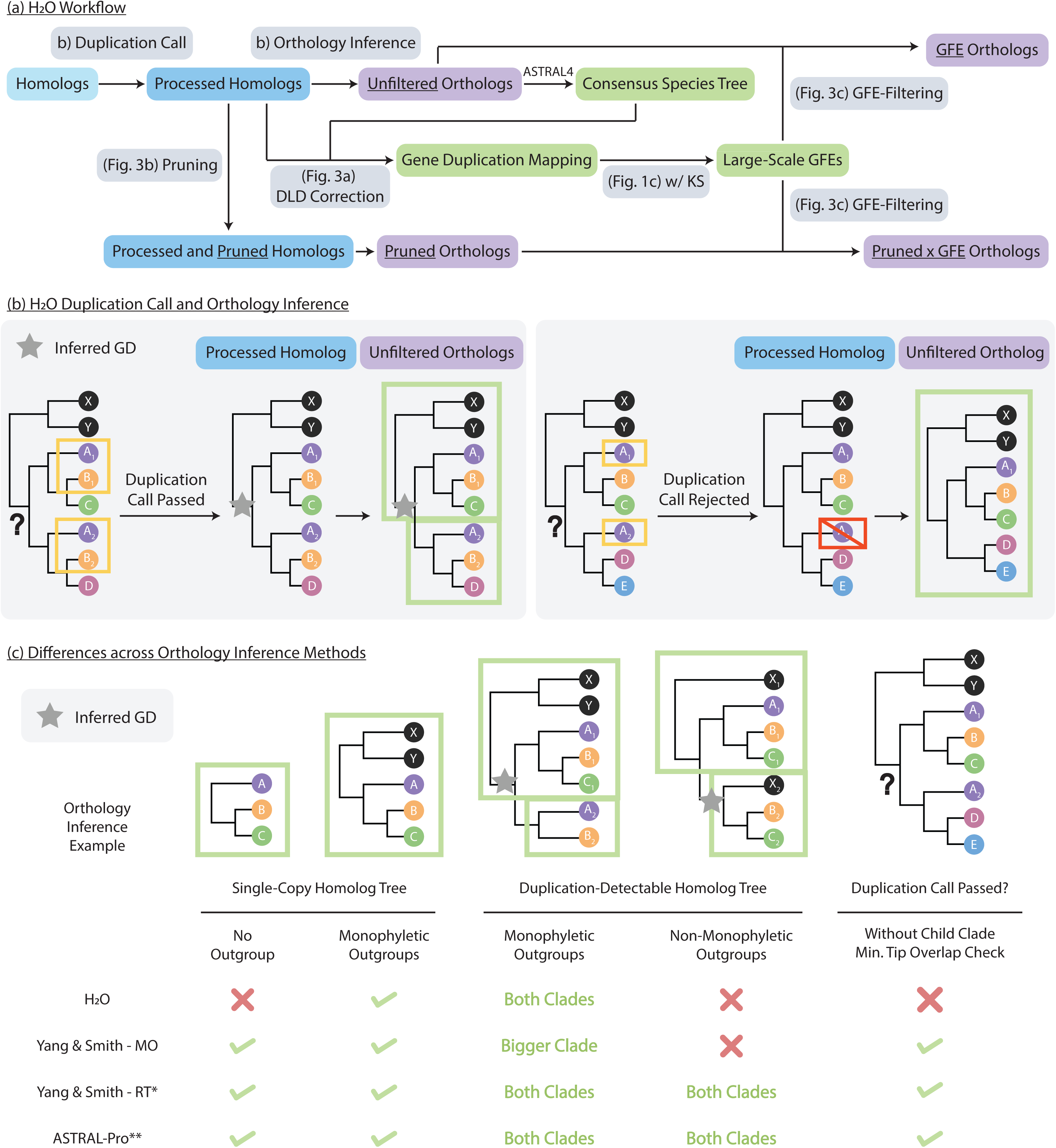
Schematics for *H_2_O*’s workflow and comparison to other ortholog inference methods. GD is short for gene duplication. Colored circles are gene sequences representing species A-E; black circles represent outgroups; subscript numbers represent duplicated gene copies. Unless otherwise specified, “homologs” and “orthologs” refer to homolog and ortholog gene trees. (a) The workflow of *H_2_O*: Homolog trees in blue (input homologs – light; processed homologs - dark); Orthologs in purple; Approaches explained in figures in gray; others in green. (b) *H_2_O* standard duplication call with child clade minimum tip overlap check and orthology inference. Question mark points to the node pending duplication call. Green rectangles enclose inferred orthologs. (c) Differences in orthology inference in all four approaches compared in this study. *RT does not output any outgroup taxa in inferred orthologs. **ASTRAL-Pro does not root homologs nor output any orthologs. This is based on how it identifies duplication nodes and which relationships are included in its consensus species tree estimation. Alt text: The figure comprises three panels illustrating H2O’s workflow and comparison to other methods. In panel a, H2O’s workflow is outlined in a flowchart, using colored rectangular boxes and directional arrows. The sequence begins with input ‘Homologs’, which progress sequentially to ‘Processed Homologs’, ‘Unfiltered Orthologs’, and ‘Consensus Species Tree’. The processes leading to processed homologs and unfiltered orthologs are detailed in panel b. Then, ‘Processed Homologs’ and ‘Consensus Species Tree’ are used in “Gene Duplication Mapping” with optional DLD correction (detailed in Figure 3a) to identify large-scale GFEs using the approaches in Figure 1C. ‘Processed Homologs’ also go through pruning (detailed in Figure 3b) to create ‘Processed and Pruned Homologs’ and ‘Pruned Orthologs’. ‘Unfiltered Orthologs’ go through GFE-filtering (detailed in Figure 3c), yielding ‘GFE Orthologs’; ‘Pruned Orthologs’ also go through GFE-filtering, yielding ‘Pruned x GFE Orthologs’. Panel b first shows a homolog tree with topology ((X,Y),((A_1_, (B_1_, C)), (A_2_,(B_2_, D)))), X and Y are outgroups. Node ((A_1_, (B_1_, C)), (A_2_,(B_2_, D))) is the node of interest. Because the two child clades have two tips overlapping (A and B), the node passed the duplication call, and two unfiltered orthologs are inferred: ((X,Y),((A_1_, (B_1_, C))) and (A_2_,(B_2_, D)). Next, when the homolog tree is ((X,Y),((A_1_, (B, C)), (A_2_,(D, E)))). The two child clades only have one tip overlap, which does not pass the duplication call, A_2_ is removed and the resulting ortholog is ((X,Y),((A_1_, (B, C)),(D, E))).

### Standard Orthology Inference

*H_2_O* first roots the homolog trees and removes trees with polyphyletic outgroups. Next, gene duplications are identified within the rooted homologous trees by comparing the child clades of each node (Figure 2b). If the two child clades of a node share at least two taxa and at least 10% of the taxa in the larger child clade, the node is recognized as a duplication node. Overlapping tips that cannot define duplication nodes are removed, and the resulting homolog trees are referred to as “processed homolog trees”. Then, the trees are partitioned into orthologous groups at each duplication node. The set of ortholog trees produced by this standard approach is referred to as “unfiltered orthologs”, as they do not undergo any special handling. Next, a consensus species tree is generated using ASTRAL4 (Tabatabaee et al. 2023; Zhang et al. 2025), though any external tool can be used here. The consensus species tree is further rooted with phyx v1.2 program pxrr (Brown et al. 2017), and gene tree discordance analysis is conducted using bellerophon (Smith et al. 2020; https://git.sr.ht/∼hms/bellerophon).

*H_2_O*’s inference of orthology from homology trees is inspired by the Rooted Ingroups (RT) and Monophyletic Outgroups (MO) methods in Yang and Smith (2014), but it is not identical to either approach (Figure 2c). *H_2_O* excludes all homolog trees with polyphyletic or no outgroups, whereas Yang and Smith (2014) RT and MO include one-to-one orthologs without outgroups. Additionally, as mentioned above, *H_2_O* has a child-clade minimum tip-overlap check for duplication node calls; a node is not recognized as a duplication node if the tip overlap between the two child clades is below the limit. Neither Yang and Smith (2014) nor ASTRAL-Pro3 (Zhang et al. 2025) includes this check, which can alter orthology inference outcomes across methods. Yang and Smith (2014) do have a minimum ortholog clade size check, but this removes small ortholog clades after the duplication node call and the dissection of the homolog tree into orthologs. ASTRAL-Pro3 (Zhang et al. 2025) does not produce ortholog trees; it roots trees in the way that minimizes the number of gene duplications and losses (often not rooting on the outgroups) without limits on minimal duplication clade size.

### Gene-duplication mapping and Large-Scale GFE Identification

From the processed homolog trees, *H_2_O* maps inferred gene duplications onto the consensus species tree with a slightly more conservative approach than PhyParts v0.0.1 (Smith et al. 2015). PhyParts (Smith et al. 2015) requires at least two tips shared by the two child clades of a node for it to qualify as a duplication node. Then, PhyParts uses the union of the descendant taxa of the focal node to map the duplication on the consensus species tree. *H_2_O* uses the same minimum tip overlap check and additionally requires that the two child clades of a node share at least 10% of the taxa in the larger child clade, to account for expression and sequencing quality differences in transcriptomes. Further, to avoid false phylogenetic placements of duplications caused by gene-tree discordance, *H_2_O* checks whether both the ingroups (descendant taxa) and outgroups (non-descendant taxa) of each duplication node in each homolog tree were congruent with those of the node in the consensus species tree. “Congruent” means the list of unique ingroup/outgroup taxa for the consensus species tree node contains all of that for the duplication node in the homolog tree. If the duplication node is incongruent with the consensus species tree, it cannot be mapped onto the consensus species tree and is therefore not recorded in the final gene duplication count for the given consensus species tree. This procedure reduces the effect of hybridization and ILS on gene duplication mapping. Each duplication is mapped onto the shallowest (furthest from the root) congruent consensus species tree node.

Additionally, *H_2_O* employs an optional duplicate-lineage discordance (DLD) correction for gene duplication mapping (Figure 3a). Duplicate-lineage discordance refers to the scenario in which the two child clades of the duplication node disagree with the duplication mapping location on the consensus species tree. DLD can often result from gene loss, ILS, hybridization, expression differences, and sequencing quality. Most sources of DLD usually do not significantly change the phylogenomic gene duplication counts for large-scale GFE events. However, DLD due to allopolyploidy can and may lead to mismatches between the Ks distribution and phylogenomic gene-duplication inference (see hybridization scenario in Figure 3a). Many detected large-scale GFEs are associated with hybridization (Wang et al. 2019a; Stull et al. 2023; Gaynor et al. unpublished data), and discrepancies between Ks distribution and phylogenomic gene- duplication mapping often center on two neighboring nested nodes (Yang et al. 2018; Feng et al. 2024). The DLD correction tests whether disagreement between Ks distributions and phylogenomics reflects consistent discordance among duplicated lineages. Therefore, we consider both gene duplication mapping results with and without DLD correction during large- scale GFE determination. DLD correction does not directly assess ILS or hybridization.

**Figure 3.**
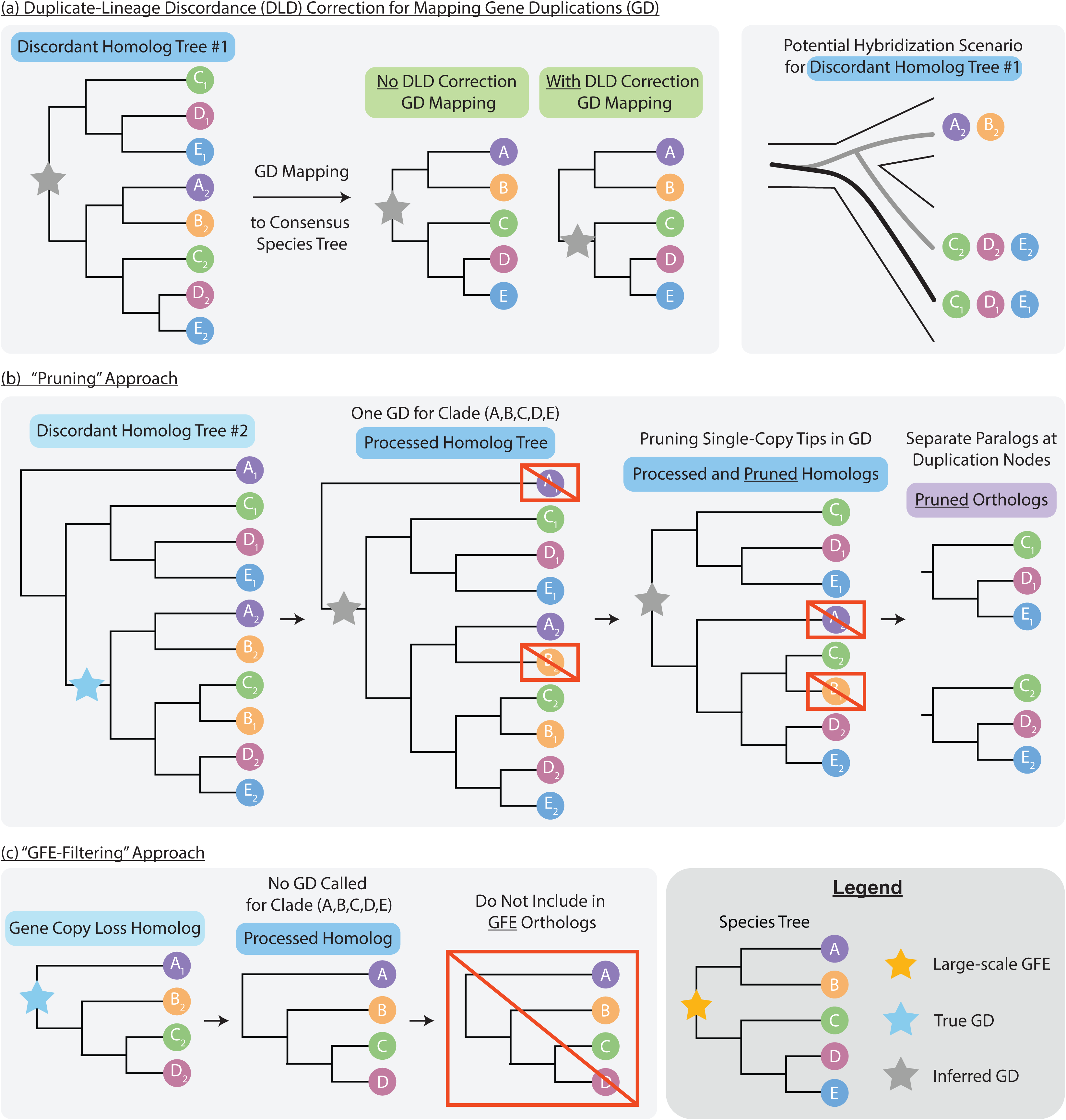
*H_2_O*’s special approaches to mitigate systematic errors in phylogenetic gene duplication and ortholog inference. GD is short for gene duplication. Colored circles are gene sequences representing species A-E; subscript numbers represent duplicated gene copies. Unless otherwise specified, “homologs” and “orthologs” refer to homolog and ortholog gene trees. (a) Optional duplicate-lineage discordance (DLD) correction for phylogenetic GD mapping. The right panel shows a potential hybridization scenario that can lead to the relationships in discordant homolog tree #1 when hybridization occurred alongside gene duplication. (b) Schematic of the “pruning” approach. (c) Schematic of the “GFE-filtering” approach. Alt text: The legend shows a consensus species tree of ((A, B), (C, (D, E))) and a large-scale GFE at the root of the tree. Panel a starts with ‘Discordant Homolog Tree #1’ of topology ((C_1_, (D_1_, E_1_)), ((A_2_, B_2_), (C_2_, (D_2_, E_2_)))). Without DLD correction, the gene duplication is mapped to the root of ((A, B), (C, (D, E))) in the consensus species tree; with DLD correction, it is mapped to the root of (C, (D, E)). Panel a also shows a lineage splitting into (A, B) and (C, (D, E). The lineage leading to (A, B) hybridized with the lineage leading to (C, (D, E) while retaining both gene copies in the (C, (D, E) lineage, which can lead to the relationships in ‘Discordant Homolog Tree #1’. Panel b starts with the input ‘Discordant Homolog Tree #2’ of topology (A_1_, ((C_1_, (D_1_, E_1_)), ((A_2_, B_2_), ((C_2_, B_1_), (D_2_, E_2_))))). The true gene duplication is at the root of ((A, B), (C, (D, E))). When the homolog tree is processed, A_1_ and B_1_ are removed because they cannot define a duplication node, and a gene duplication is inferred at the root of the homolog tree. In pruning, single-copy A_2_ and B_2_ are removed, resulting in two pruned orthologs (C_1_, (D_1_, E_1_)) and (C_2_, (D_2_, E_2_)). Panel c starts with an input ‘Gene Copy Loss Homolog’ of (A_1_, (B_2_, (D_2_, E_2_))), and the ‘Processed Homolog’ and the ortholog tree have the same topology. This ortholog is not included in GFE orthologs.

To detect large-scale GFE events, we highlighted the Ks values of each gene-duplication- clustered node in the Ks distributions (node-partitioned Ks distributions, Figure 1c). To identify paralog Ks duplicated at each node, *H_2_O* outputs a list of paralog gene pairs and their corresponding duplication nodes in the consensus species tree during the gene-duplication mapping process. We generated Ks values for all paralogous gene pairs using homologous alignments. Here, we used a modified Ks calculation protocol following Yang et al. (2015), with Nei−Gojobori correction for multiple substitutions. We modified synonymous_calc.py (https://github.com/tanghaibao/bio-pipeline/blob/main/synonymous_calculation/synonymous_calc.py) to calculate Ks in an all-by-all pairwise approach for each species in each homolog cluster, instead of only calculating Ks for non-overlapping adjacent pairs. Subsequently, we isolated Ks values for paralogs duplicated from each node of interest, i.e., those with a high number of mapped gene duplications and/or corresponding to a putative large-scale GFE event suggested by previous studies. To recognize putative WGDs, we also obtained median chromosome counts from the Chromosome Counts Database v1.66.5 (Rice et al. 2015).

Considering the node-partitioned Ks distributions, phylogenetically mapped gene duplications, and chromosome counts together, we placed large-scale GFE and putative WGDs on the consensus species tree. If placement was still uncertain, we calculated Ks for orthologous gene pairs from species diverged at nearby nodes with the same Python script and compared them with those of paralogous gene pairs to verify gene duplication timing. We acquired the ortholog gene pairs through a custom script. Ortholog Ks is a rough reflection of the timing of speciation, and paralog Ks is an approximation for the timing of gene duplication.

### Homolog and Ortholog Special Filtering Methods

*H_2_O* employs two special data filtering methods for orthology inference, based on two hypotheses about potential sources of hidden paralogy and systematic errors that may contribute to the apparent association between large-scale GFE and gene tree discordance.

First, because many ancient WGDs are also products of ancient hybridization (allopolyploidy) (Wang et al. 2019a; Stull et al. 2023; Gaynor et al. unpublished data), the interplay among gene duplication, loss, and hybridization can further confound orthology inference. Under these circumstances, various methods may recover different orthology relationships from the same homologous gene family. Second, diploidization following WGD leads to extensive loss of duplicate genes (Li et al. 2021; Wang et al. 2021). While some gene families retain the duplicate copies, others return to single-copy status. As duplicated gene loss is lineage-specific and non- random, we cannot assume all single-copy genes are orthologs if the lineage experienced large- scale GFEs (Carretero-Paulet and Fares 2012; Paterson et al. 2012; Moghe et al. 2014; Feng et al. 2024). Here, “duplicate gene loss” refers broadly to the absence of one or more gene copies from a transcriptomic dataset for the sampled species. The loss can result from true gene loss, expression differences, or sequencing and assembly errors. Nevertheless, current phylogenetic orthology inference methods assume single-copy gene trees represent valid orthologs (Marcet- Houben et al. 2011; Yang and Smith 2014; Ballesteros and Hormiga 2016; Zhang et al. 2025).

This hidden paralogy may lead to erroneous orthology inference and spurious gene tree conflict – discordance driven by systematic error rather than biological processes.

To address these hypotheses, we introduce two specialized data filtering approaches for orthology inference to circumvent hidden paralogy and reduce systematic errors: (1) the “pruning” approach (Figure 3b) and (2) the “GFE” approach (Figure 3c). As mentioned above, there are two potential sources of systematic errors from the uncertainty of true orthology: when (1) ILS/hybridization coincides with large-scale GFEs, and (2) only one of the duplicated gene copies is retained for every descending species following large-scale GFE events. To resolve the first problem, for each gene duplication, *H_2_O* removes tips that no longer retain both gene copies generated by that duplication, before breaking homologous trees into orthologs (the “pruning” approach; Figure 3b). With pruning, we are more certain about gene orthology, as all remaining tips retained both copies in duplicated clades. However, this approach does not test for ILS or hybridization. It could remove hidden paralogs, but it could also remove orthologs. The modified homolog and ortholog trees are subsequently referred to as “pruned homolog/ortholog trees”. To resolve the second issue, *H_2_O* constructs a subset of ortholog trees inferred from homolog trees that retain duplicated gene copies generated by each selected large-scale GFE event associated with gene tree discordance (the “GFE-filtering” approach; Figure 3c). Similarly, the GFE-filtering does not test for hidden paralogy. It could remove hidden paralogs and also single-copy orthologs. These filtered ortholog trees are subsequently referred to as “GFE ortholog trees”. When both approaches are applied, the set of ortholog trees is referred to as “pruned x GFE trees”. For each large-scale GFE that exhibits gene tree discordance, a “GFE” and a “pruned x GFE ortholog tree” set are generated for each event. Thus, one dataset can have more than one set of “GFE” and “pruned x GFE” ortholog trees.

### Simulation

We evaluated *H_2_O*’s orthology inference outcome against Yang and Smith (2014) RT and MO and ASTRAL-Pro3 (Zhang et al. 2025) using a simulated homologous tree dataset. We conducted the tree simulation with SimPhy (Mallo et al. 2016) and custom scripts. The simulated homologous trees mimicked scenarios in which a large-scale GFE event occurred just before the crown node of a clade, with different levels of ILS around this node. We tested three levels of ILS: high, medium, and none. We aimed to resemble empirical-level gene tree discordance associated with large-scale GFEs using high-ILS simulation. The medium-ILS level was an intermediate between high-ILS and no ILS; the no-ILS simulation was a control setting. The details of tree simulations are reported in supplementary text.

After simulating the homolog gene trees, we ran the *H_2_O* pipeline on the test sets for each ILS level. For each test homolog tree set, *H_2_O* produced four sets of ortholog gene trees: (1) unfiltered orthologs with the standard approach, (2) pruned orthologs, (3) GFE orthologs, and (4) pruned x GFE orthologs. Then, we ran ASTRAL-Pro3 (Zhang et al. 2020; Tabatabaee et al. 2023; Zhang et al. 2025) and Yang and Smith (2014) on each ILS test set. ASTRAL-Pro3 yielded a consensus species tree directly from homolog trees, with more rounds to optimize the output topology. For Yang and Smith (2014), we ran the two orthology inference methods most similar to *H_2_O*, Rooted Ingroups (RT) and Monophyletic Outgroups (MO), for each clade, and each produced a set of ortholog trees. We then used ASTRAL4 (Zhang et al. 2025) to produce consensus species trees for each set of orthologs. We rooted each consensus species tree with phyx v1.2 program pxrr (Brown et al. 2017) and conducted gene tree discordance analyses with bellerophon (Smith et al. 2020; https://git.sr.ht/∼hms/bellerophon). Bellerophon is a time- efficient tool for large datasets that counts the number of conflicting and concordant ortholog gene trees with a given consensus species tree. Because bellerophon takes ortholog trees as input, and ASTRAL-Pro3 does not automatically produce ortholog trees, we ran ASTRAL-Pro3 to produce homolog trees with duplication nodes labeled. We rooted the labeled homolog trees with the outgroups, dissected them into ortholog trees with a custom script, and used them as input for bellerophon. Additionally, we ran bellerophon on the control single-copy homolog tree sets and used a custom script to evaluate orthology-inference accuracy for each method against the true orthology.

### Empirical Validation

We tested *H_2_O* on three empirical phylotranscriptomic datasets: Ericales (ERIC; Carruthers et al. 2024), and two Caryophyllales subclades, Caryophyllaceae (CARY; Feng et al. 2024) and the “Phytolaccoid clade” + Aizoaceae (PHYT; Yang et al. 2018). We obtained homologous phylogenetic trees and corresponding sequence alignments from the source publications. We also updated the species names for all datasets using the World Checklist of Vascular Plants (WCVP, updated January 2026; POWO 2026; see Table S1). The homology inference in all three studies was conducted with the Yang and Smith (2014) pipeline. For orthology inference, Yang et al. (2018) exclusively used single-copy homolog trees as the orthologs; Carruthers et al. (2024) and Feng et al. (2024) used Yang and Smith (2014) RT. We conducted further cleaning for some homologs (details see supplementary materials).

We applied *H_2_O*’s workflow to all three transcriptomic datasets. We analyzed only GFEs shared by at least two species, because multiple transcripts sequenced within a single species may not indicate real gene duplications. To explore the relationship between Ks values and life history, we obtained life history data for CARY from Feng et al. (2024) and for the other two clades from Smith et al. (2026). We grouped annual and perennial herbaceous plants as herbaceous; shrubs, trees, and lianas as woody. We selected large-scale GFEs shared by a relatively large group of species in each dataset and conducted independent two-sample t-tests on the Ks peak means for the event between herbaceous and woody species. We further estimated the phylogenetic signal (Pagel’s λ) of Ks peak means and conducted phylogenetic ANOVA tests comparing those of herbaceous and woody species with the R package phytools (Revell 2024).

### Evaluating H_2_O’s Orthology Inference Outcome on Empirical Datasets

We employed *H_2_O*’s workflow on the three empirical phylotranscriptomic datasets in the same way as the simulated dataset, which resulted in four sets of ortholog trees: (1) unfiltered orthologs with the standard approach, (2) pruned orthologs, (3) GFE orthologs, and (4) pruned x GFE orthologs. For comparison, we also ran ASTRAL-Pro3 (Zhang et al. 2025) and Yang and Smith (2014) RT and MO, and conducted gene tree discordance analyses the same way as with the simulated dataset.

We defined gene tree discordance as the proportion of conflicting gene trees relative to the total number of conflicting and concordant gene trees, compared with the consensus species tree. To compare discordance across methods, we selected large-scale GFE events associated with high levels of gene tree discordance and distinguished the neighboring descendant nodes of these events (i.e., GFE-associated nodes) from other nodes in the trees. Using the *H_2_O* “pruned x GFE” method as a baseline, we calculated the difference in discordance between each method and the baseline for each GFE-associated node. Then we conducted paired two- sample t-tests to assess whether the differences were positive. Lastly, we obtained the number of ortholog trees and the average number of tips per tree for each method. Using *H_2_O* “unfiltered” ortholog trees as a benchmark, we quantified the proportional change in these two numbers for each method relative to the benchmark.

### Examining Trends in Gene Family Contractions

First, we explored trends in gene copy losses after each major large-scale GFE event (i.e., gene family contractions). We consider large-scale GFE events “major” when they are shared by many species and are associated with relatively large numbers of gene duplications within the clade. We mapped the gene copy losses with *H_2_O*. To map gene loss for the selected GFE events, *H_2_O* obtained the list of taxa that were missing in each orthologous gene copy for each gene duplication associated with the event and located the most parsimonious node(s) where the gene copy was lost. Gene copy losses were mapped only when all descendant taxa of that node were missing that gene copy in the homolog tree. Single-tip losses were disregarded because they are more likely to be a consequence of tissue-specific gene expression and/or sequencing than of gene copy loss.

Second, we explored the relationship between gene family contractions and molecular branch length. The consensus species trees used here are those inferred from “unfiltered” orthologs.

To obtain molecular branch lengths, we re-estimated the branch lengths of the consensus species tree, detailed in supplementary text. We applied linear regression models to the number of gene copy losses after each large-GFE event, as a function of the molecular branch length leading to the node to which the loss is mapped. To categorize gene family contractions, we distinguished named clade nodes (family, tribe, genus) from other nodes in the consensus species tree. We calculated the proportion of gene family contractions as the number of gene copy losses divided by the total number of gene duplications in the large-scale GFE. Then, for each large-scale GFE event, we performed independent two-sample t-tests comparing the proportion of gene family contractions mapped right before named clades with that of those not mapped before named clades, with and without correcting for molecular branch lengths.

We also conducted independent two-sample t-tests comparing molecular branch lengths leading to named clades with those leading to other nodes.

We used Cursor, GitHub Copilot Auto AI models, and Claude Sonnet 5.5 to assist with writing and optimizing code. All scripts used in this study were manually reviewed and tested.

## Results

### H_2_O Effectively Filtered Hidden Paralogs in Simulated Datasets

We compared the inferred ortholog tree topology of the seven methods to the true simulated orthologs: *H_2_O* unfiltered, *H_2_O* pruned, *H_2_O* GFE, *H_2_O* pruned x GFE, ASTRAL-Pro3 (Zhang et al. 2025), Yang and Smith (2014) RT and MO. For the high-ILS case, in which the proportion of gene tree discordance is the closest to what we observe in empirical datasets, the proportion of paralogous relationships reported and gene tree discordance were similar among *H_2_O* unfiltered, ASTRAL-Pro3 (Zhang et al. 2025), Yang and Smith (2014) RT and MO (Figure 4). These paralogous relationships were mostly inferred from single-copy (hidden paralog), duplication- detectable (monophyletic), and duplication-detectable (hidden paralog) homolog trees. Single- copy (hidden paralog) homolog trees had the highest counts because they made up 30% of the total simulation homolog trees, and no orthology inference method could differentiate paralogs in a single-copy homolog tree.

**Figure 4.**
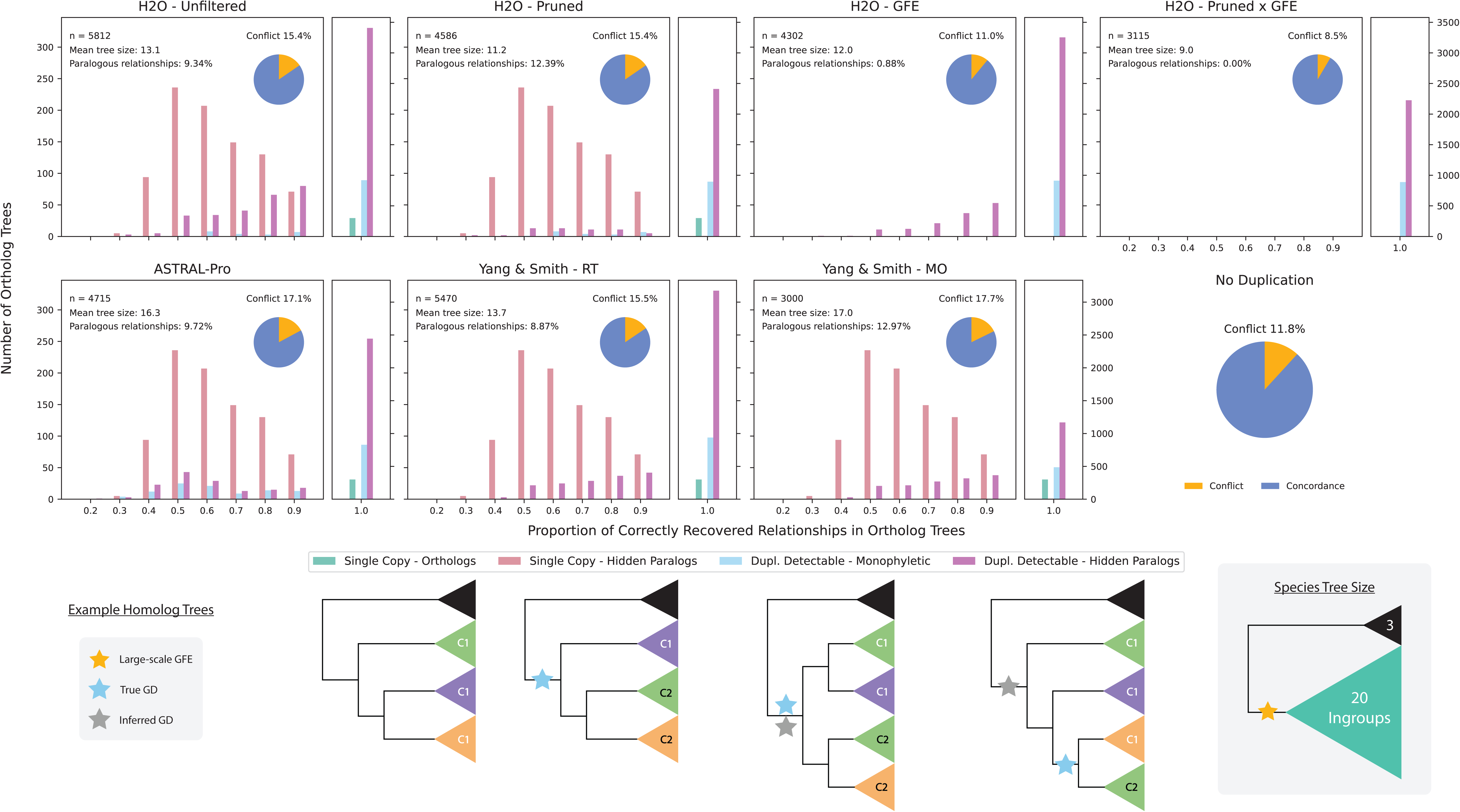
Simulation results for high-ILS setting. The top eight panels represent the simulation results for the seven sets of inferred ortholog trees and the simulated single-copy homolog control tree set (no duplication). The x-axis of the histogram represents the proportion of correctly recovered ortholog relationships in each method compared to the simulated true orthologs (1.0 means that the inferred ortholog tree was only comprised of true orthologs). For readability of the histograms in the range x < 1, each panel is divided into two subpanels: x < 1 (y-axis labeled on the left) and x = 1 (y-axis labeled on the right). “n” is the number of ortholog trees inferred; mean tree size is the mean number of tips per tree, including outgroups; paralogous relationships are proportional to the total number of inferred relationships. The pie charts report gene tree discordance summed across all nodes in the consensus species tree. The colors of the histogram represent the type of simulated homolog tree, with example topologies below. C1 and C2 refer to gene copy 1 and copy 2; triangle color indicates different ingroup clades. Alt text: The histograms of seven sets of ortholog trees show that H2O unfiltered, ASTRAL-pro, and Yang and Smith (2014) RT and MO report similar amounts of paralogous relationships inferred from single-copy (hidden paralog), duplication-detectable (monophyletic), and duplication-detectable (hidden paralog) homolog trees. H2O “pruned” filter removed paralogous relationships inferred from duplication-detectable (hidden paralog) homolog trees. H2O “GFE” filter removed paralogous relationships inferred from single-copy (hidden paralog) homologs but also removed single-copy (ortholog) trees. H2O “pruned x GFE” removed all paralogous relationships reported in the unfiltered orthologs and resulted in less gene tree discordance compared to the single-copy control homolog tree set.

*H_2_O* “pruned” filter removed most paralogous relationships inferred from duplication- detectable (hidden paralog) homolog trees, but it also removed correctly inferred relationships from them, thus resulting in a higher proportion of reported paralogous relationships. *H_2_O* “GFE” filter removed all single-copy gene trees, eliminating paralogous relationships inferred from single-copy (hidden paralog) homologs but also discarded single-copy (ortholog) trees.

Although *H_2_O* “pruned x GFE” removed many inferred ortholog trees and tips from the unfiltered orthologs, it reported no paralogous relationships and produced less gene tree discordance than the single-copy control homolog tree set, which represented the proportion of gene tree discordance only produced by ILS. Results from median-ILS level and no-ILS were similar, as shown in Figure S1.

### H_2_O Reduced Uncertainty in Large-Scale GFE Detection

All three phylogenomic datasets contain multiple putative ancient WGDs (i.e., large-scale GFEs) identified in their original studies; some remain uncertain because Ks distribution peaks did not match gene duplication count mappings. We re-evaluated event identifications in each dataset by highlighting the Ks values of paralogs duplicated at nodes with gene duplication clustering. Most phylogenetically inferred paralogs have Ks values of less than two, as they were in gene families well represented within their own dataset and thus had short molecular distances between gene copies duplicated within the clade. Events that are only ancestral to one species are not considered in this study. We address the main results here while the specifics are discussed in supplementary materials.

Caryophyllaceae (CARY) is a strong example showing that node-partitioned Ks distributions and DLD corrections reduced uncertainty in large-scale GFE detection (Figure 5; Figures S2-S9). We observed clear highlighted peaks in the node-partitioned Ks distributions for CARY1-CARY3 (Figure 3b) and clear differentiation in highlighted Ks peaks for different large-scale GFE events within the same species (Figure 3c, *Schiedea membranacea* and *Colobanthus subulatus*).

**Figure 5.**
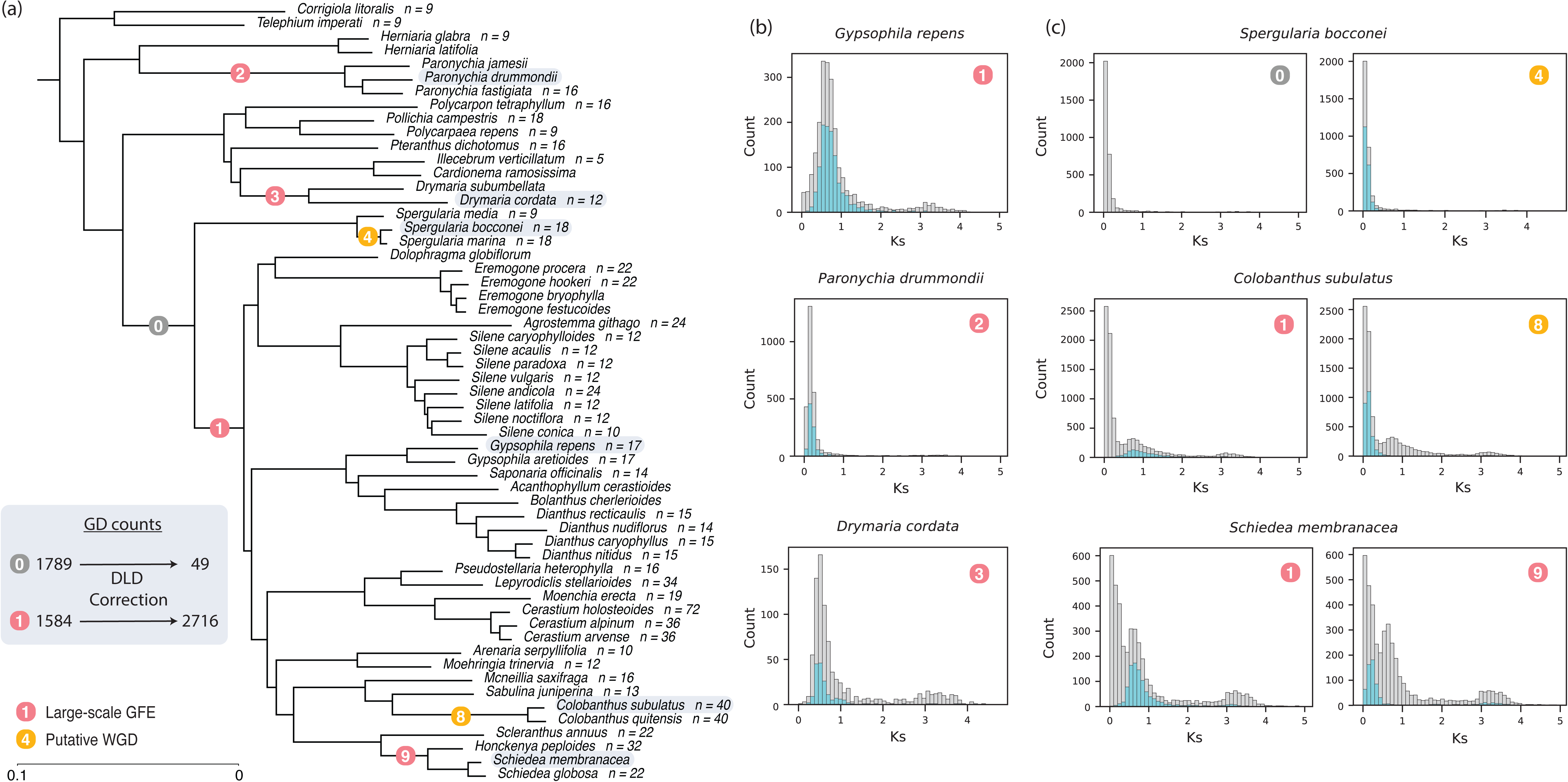
Highlighting phylogenomically inferred gene duplications in Ks distributions validated large-scale GFE detection in Caryophyllaceae (CARY). (a) Large-scale GFE/WGD events labeled on the consensus species tree with molecular branch length (substitutions per site). Colored circles are large-scale GFE/WGD event numbers for CARY1-6 and 8-9. Only selected cases of GFEs are plotted, so the numbers are not consecutive. The gray circle is not an event (node 0), although many gene duplications mapped to it. GD is short for gene duplication. Median chromosome counts are labeled to the right of species names. Putative WGDs are supported by increases in the base chromosome numbers. Species included in (b) and (c) are highlighted in gray. (b) and (c) are examples of node-partitioned Ks distributions. Gray bars are Ks for all paralog gene pairs; blue bars are Ks for paralog gene pairs diverged from the event labeled in the plot (with no DLD correction). (b) is for species that experienced a single event. (c) compares node-partitioned Ks for gene duplications from different events. Alt text: Panel a shows a consensus species tree of Caryophyllaceae, where terminal branches are annotated with median chromosome counts. Panels b and c are two sets of histograms displaying the node-partitioned Ks distributions for paralog gene pairs, with clear peaks for each large-scale GFE event. One species is chosen as an example for each event. Additionally, panel C shows that Ks peaks from different GFE events have distinct mean values within each species.

Without DLD correction, gene duplication mapping does not fully match standard Ks distributions for CARY1. All descendant species of node 0 (Figure 5a) except *Spergularia media* had a standard Ks peak with similar means (Figure S3); both node 0 and CARY1 had a large number of gene duplications mapped to them without DLD correction (Figure 5a). However, DLD correction resolved this discrepancy, indicating that most gene families with a gene duplication mapped to node 0 were single-copy in *Spergularia*. This is evident in the node- partitioned Ks distributions for *Spergularia bocconei* (Figure 3b), as no Ks peak was highlighted for node 0.

We resolved a similar discrepancy for a putative WGD, ERIC1, in Ericales (ERIC) with DLD correction and node-partitioned Ks distributions (Figures S10-S14). Our determination is consistent with synteny analysis results in Zhang et al. (2020), while at odds with Carruthers et al. (2024). *H_2_O*’s conservative gene-duplication mapping cleaned up the scattered gene duplications mapped along the backbone of Ericales in Carruthers et al. (2024) by checking concordance between duplication nodes in the homolog gene tree and nodes in the consensus species tree (Figure S10). This also facilitated large-scale GFE detection.

For the “Phytolaccoid clade” and Aizoaceae (PHYT), our large-scale GFE determination (Figures S15-S19) is consistent with those reported in Yang et al. (2018). Notably, there was a significant deviation in Ks peak means between woody and herbaceous descendant species of PHYT1 (Figure S17 and Table 1; *p* = 5.93 × 10^−16^). We conducted the same statistical test for ERIC1 and CARY1; the former was significant, and the latter was not (Table 1). However, these results should be interpreted with caution due to the strong phylogenetic signal of Ks (Table 1). Given these strong phylogenetic signals, we conducted phylogenetic ANOVA tests, and none were statistically significant. The phylogenetic ANOVA may fail to reject the null hypothesis due to the small number of life-history shifts (e.g., only one in PHYT1) and small sample sizes.

**Table 1.**
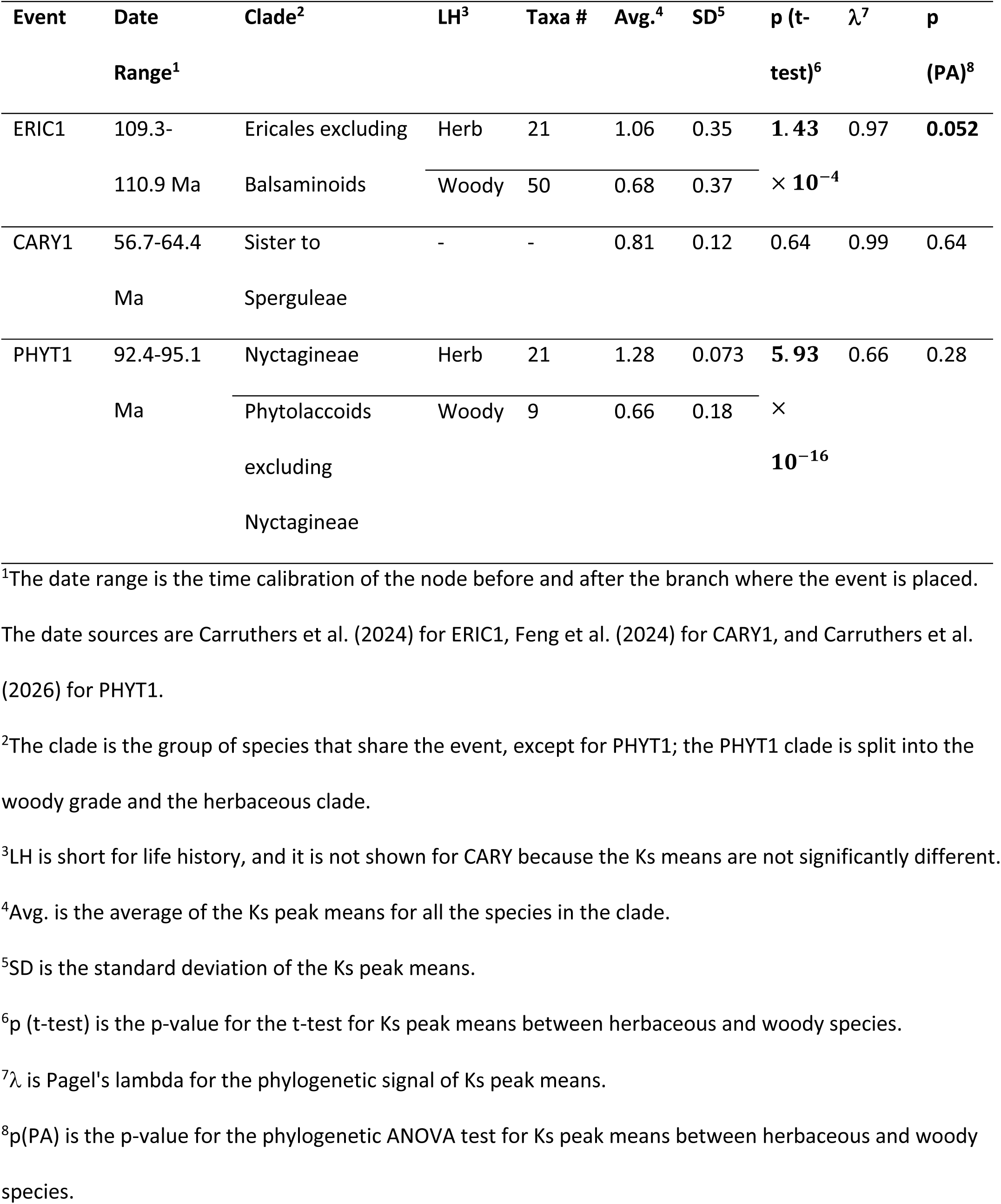
Time ranges and averages of Ks peak means of major large-scale GFE events, compared between herbaceous and woody species.

| **Event** | **Date Range<sup>1</sup>** | **Clade<sup>2</sup>** | **LH<sup>3</sup>** | **Taxa #** | **Avg.<sup>4</sup>** | **SD<sup>5</sup>** | **p (t-test)<sup>6</sup>** | **λ<sup>7</sup>** | **p (PA)<sup>8</sup>** |
|---|---|---|---|---|---|---|---|---|---|
| ERIC1 | 109.3-110.9 Ma | Ericales excluding Balsaminoids | Herb | 21 | 1.06 | 0.35 | **1.43** × **10<sup>−4</sup>** | 0.97 | **0.052** |
|  |  |  | Woody | 50 | 0.68 | 0.37 |  |  |  |
| CARY1 | 56.7-64.4 Ma | Sister to Sperguleae | - | - | 0.81 | 0.12 | 0.64 | 0.99 | 0.64 |
| PHYT1 | 92.4-95.1 Ma | Nyctagineae | Herb | 21 | 1.28 | 0.073 | **5.93** × **10<sup>−16</sup>** | 0.66 | 0.28 |
|  |  | Phytolaccoids excluding Nyctagineae | Woody | 9 | 0.66 | 0.18 |  |  |  |

<sup>1</sup>The date range is the time calibration of the node before and after the branch where the event is placed. The date sources are Carruthers et al. (2024) for ERIC1, Feng et al. (2024) for CARY1, and Carruthers et al. (2026) for PHYT1.
<sup>2</sup>The clade is the group of species that share the event, except for PHYT1; the PHYT1 clade is split into the woody grade and the herbaceous clade.
<sup>3</sup>LH is short for life history, and it is not shown for CARY because the Ks means are not significantly different.
<sup>4</sup>Avg. is the average of the Ks peak means for all the species in the clade.
<sup>5</sup>SD is the standard deviation of the Ks peak means.
<sup>6</sup>p (t-test) is the p-value for the t-test for Ks peak means between herbaceous and woody species.
<sup>7</sup>λ is Pagel's lambda for the phylogenetic signal of Ks peak means.
<sup>8</sup>p(PA) is the p-value for the phylogenetic ANOVA test for Ks peak means between herbaceous and woody species.

### H_2_O Reduced Gene Tree Discordance Associated with GFEs in Empirical Datasets

For each of the three phylotranscriptomic datasets, the *H_2_O* pipeline employed two approaches, “pruning” and “GFE”, to reduce systematic error in orthology inference by avoiding hidden paralogy. This produced four sets of ortholog trees: (1) “unfiltered”, (2) “pruned”, (3) “GFE”, and (4) “pruned x GFE”. For the GFE approach, we selected six large-scale GFE events associated with gene tree discordance in their dataset: ERIC1 (Ericales excluding Balsaminoids), CARY1 (sister to Sperguleae), CARY6w (*Silene*), CARY7w (*Cerastium*), PHYT1 (Phytolaccoids), and PHYT2 (Nyctagineae). For each of the selected GFE events, a set of “GFE” and “pruned x GFE” ortholog trees was produced. The tree topologies and gene tree discordances for each method and clade are shown in Figures S20-S25.

Across all methods, mean discordances mostly did not differ for the GFE-associated nodes except for “pruned x GFE” (Figure 6a), similar to simulation results. For those that do differ, ASTRAL-Pro’s mean discordance was statistically significantly lower than MO (*p* = 0.019) and unfiltered (*p* = 0.001). Despite substantial gene tree discordance in ASTRAL-Pro3 results, posterior probability (PP) values are mostly 1.0 across all three datasets, with a few exceptions: ERIC has five nodes at ∼0.99 (Figure S24), and PHYT has two nodes at 0.81 and 0.94 (Figure S25).

**Figure 6.**
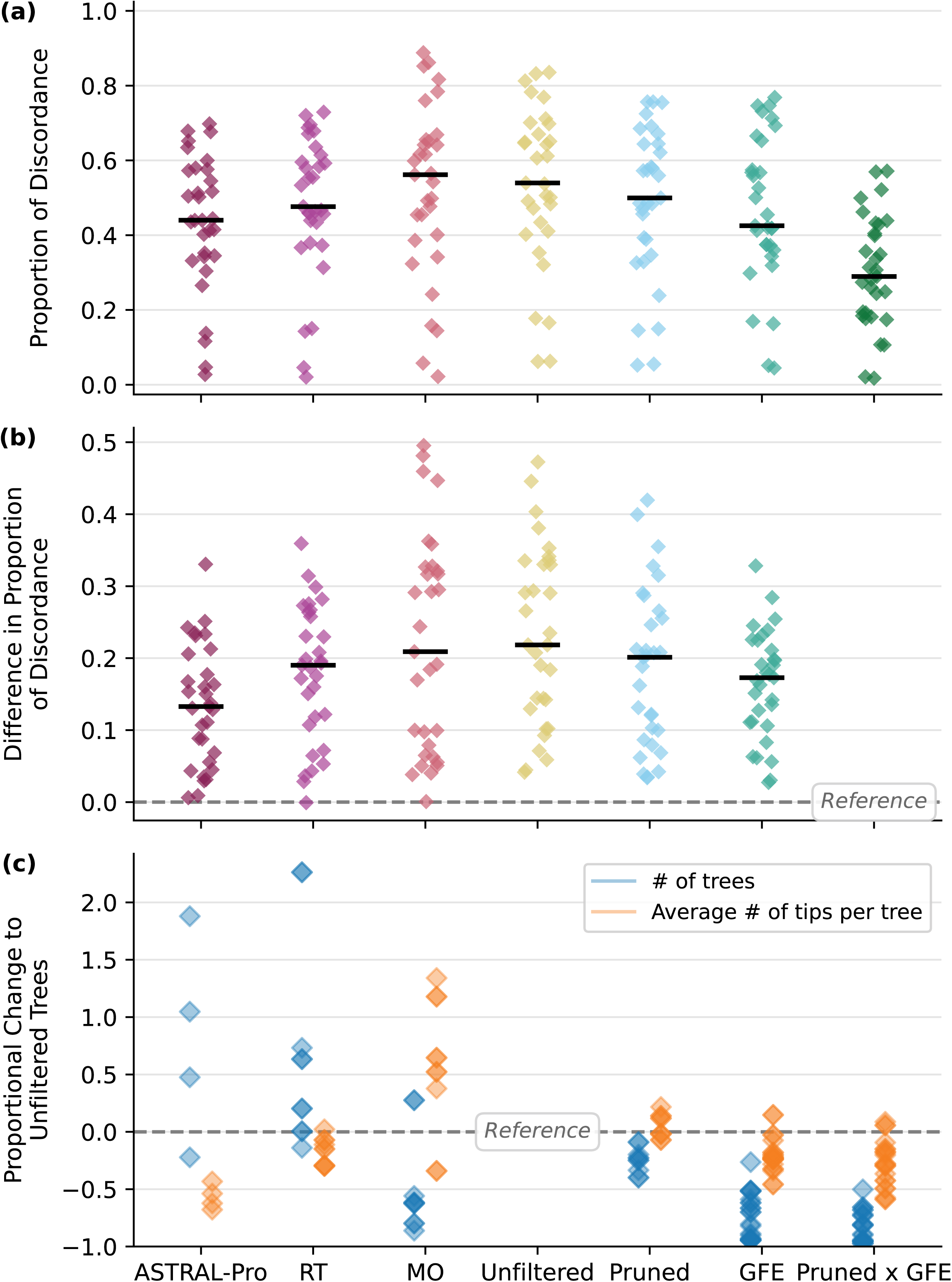
Comparing orthology inference methods within *H_2_O* and with ASTRAL-Pro and Yang and Smith (2014) (MO and RT) across all three empirical datasets. (a) Proportion of gene tree discordance in GFE-associated nodes in the consensus species tree across seven methods. (b) Proportion of discordance in each corresponding GFE-associated node in each method minus that of “pruned x GFE”. (c) Proportional change in the number of total ortholog trees produced and the average number of tips per ortholog tree, compared to “unfiltered” orthologs. The y- axis is positive when the value is higher in the selected method. The lowest run still had about 1000 gene trees for Pruned x GFE. Specific numbers are reported in Table S2. Alt text: Three-panel composite figure presents comparative analyses of orthology inference methods using scatterplots. Panel a features scatterplots for seven methods—ASTRAL-Pro3, RT, MO, Unfiltered, Pruned, GFE, and Pruned x GFE. The y-axis displays the proportion of discordance. Pruned x GFE has slightly lower discordance compared to other methods. Panel b uses scatterplots to show that in each GFE-associated node, discordance is always equal to or higher in other methods compared to Pruned x GFE. Panel c shows that, compared to the unfiltered method, Pruned, GFE, and Pruned x GFE methods generally have lower numbers of trees; GFE and Pruned x GFE methods generally have lower average number of tips per tree.

The average gene tree discordance of “pruned x GFE” was significantly lower than that estimated under the other six methods (ASTRAL-Pro3: *p* = 1.04 × 10^−4^; RT: *p* = 1.16 × 10^−5^; MO: *p* = 3.42 × 10^−6^; “unfiltered”: *p* = 8.50 × 10^−8^; “pruned”: *p* = 2.93 × 10^−6^; “GFE”: *p* = 1.09 × 10^−7^). In each GFE-associated node selected, “pruned x GFE” orthologs were less discordant with the inferred consensus species tree compared to all other methods (Figure 6b). This reduced discordance is also accompanied by a different recovered consensus species tree topology in the Ericales backbone relationships following ERIC1 (Figure S23). As in the simulation, *H_2_O* “pruned x GFE” substantially reduced gene tree discordance by removing substantial amounts of data (Figure 6c; exact numbers in Table S2). The number of ortholog trees and the average number of tips are much lower for “pruned x GFE” compared to the baseline “unfiltered” approach, for most runs. This resulted in low resolution in some relationships far from the GFE event, such as ERIC1 (Figure S23) and CARY6w (Figure S21).

### Trends in Gene Family Contractions

To explore gene family contractions, we selected major large-scale GFE events: ERIC1 (Ericales excluding Balsaminoids), CARY1 (sister to Sperguleae), PHYT1 (Phytolaccoids), and PHYT2 (Nyctagineae). For each selected large-scale GFE, we chose different taxonomic ranks for the “named clades”: ERIC1 – family; CARY1 – tribe and genus; PHYT1 – family; PHYT2 – genus. The size of inferred gene-family contractions, measured as the number of mapped duplicate-copy absences of each node following each large-scale GFE, is weakly (0.2 < *R*^2^ < 0.5) positively correlated in ERIC1, CARY1, and PHYT1, and moderately (0.5 < *R*^2^ < 0.6) positively correlated in PHYT2 (Figure S26). Larger inferred contractions tend to map to branches leading to named clades, including tribes, families, and genera, while longer molecular branch lengths are also more likely to occur before named clades (Figure 7a and 7b). The outlier in the PHYT2 other nodes in Figure 7a is the MRCA of *Tripterocalyx* and *Abronia* (Nyctaginaceae). *Tripterocalyx* was once classified under *Abronia,* and the two genera are morphologically similar, although they do not currently define a named clade. As the sizes of gene family contractions are weakly to moderately correlated with molecular branch length in some clades, we residualized the sizes with respect to branch length. After correcting for branch length, the sizes of gene family contractions no longer differ between named clades and others (Figure 7c).

**Figure 7.**
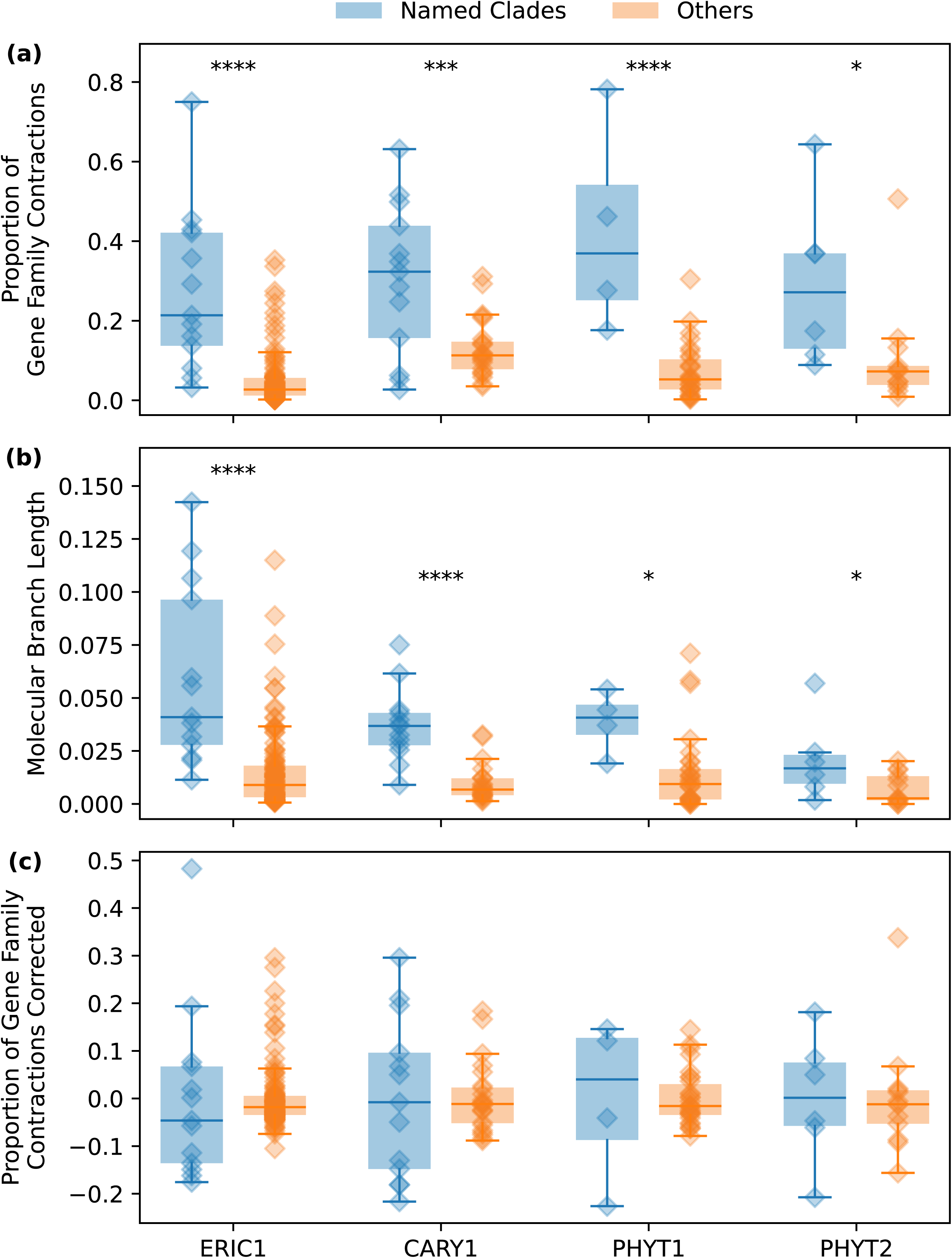
Trends in gene family contractions. (a) Boxplots for the proportion of gene family contractions between named clades and other nodes. The x-axis represents large-scale GFE events from which gene family contractions are inferred; the y-axis shows the proportion of gene family contractions relative to the number of gene duplications of the GFE event. (b) Boxplots for the molecular branch lengths directly preceding named clades and those of other nodes. (c) Boxplots for the proportion of gene family contractions corrected by molecular branch lengths between named clades and other nodes. Alt text: Panel a presents boxplots for the proportion of gene family contractions contrasting named clades and other nodes. In all four events (ERIC1, CARY1, PHYT1, PHYT2), named clades have a significantly higher proportion of gene family contractions. Panel b features boxplots of molecular branch lengths for the same events, with named clades generally exhibiting longer branch lengths. Panel c provides boxplots of the proportion of gene family contractions corrected by molecular branch length, again comparing named clades and other nodes across seven events; none of the comparisons are statistically significant.

## Discussion

### Confident Detection of Large-Scale Gene Family Expansions

Across the three phylotranscriptomic datasets of Ericales and two Caryophyllales subclades, the detected large-scale GFE events vary in the number of associated gene duplications and mostly lack both a base chromosome count increase and synteny support from existing studies.

Despite variation in the number of inferred gene duplications, many detected events exhibit consistent highlighted-Ks peaks for the corresponding paralog pairs across sampled descendant species in the node-partitioned Ks distributions. This pattern is less consistent with species- specific transcript assembly artifacts or isoform inflation, which should primarily generate idiosyncratic, lineage-specific signals and Ks values concentrated near zero. Nevertheless, transcriptomic data alone cannot eliminate all assembly-related error sources, and variation in the number of mapped duplications across events may reflect a mix of biological processes, such as diploidization and differential duplicate retention, and technical factors, including tissue-specific gene expression, transcript recovery, assembly quality, and taxon sampling. For this reason, we use the term “large-scale GFE” rather than “WGD” for events inferred primarily from transcriptomic Ks distributions and gene-tree analyses, reserving “putative WGD” for cases with additional evidence such as chromosome-number shifts or syntenic support. In many downstream comparative analyses, the immediate biological process of interest may be the history of retained gene-family expansion, regardless of whether the original source was whole- genome duplication, segmental duplication, or another large-scale duplication process.

Using node-partitioned Ks distributions that integrate phylogenetic placement and Ks values for paralogous gene pairs in a single analysis was key to confidently identifying large-scale GFEs.

Discrepancies between Ks distributions and phylogenomically inferred gene duplication confounded large-scale GFE placements in the source studies of the empirical datasets. The inconsistencies are also evident in our results, as many gene-duplication clusterings on the consensus species tree were not previously recognized as large-scale GFEs (Figures S2, S10, and S15). Bridging the two methods allowed us to determine both the timing and phylogenetic placement of GFEs (Figure 1c) and to resolve discrepancies in detecting and locating large-scale GFEs across all three datasets. In node-partitioned Ks distributions, because Ks distributions were calculated from duplicate pairs within gene families filtered by phylogenetic criteria, they are not independent of the gene-tree inferences. Instead, they provide complementary information on the relative timing of duplication events identified through phylogenomic analysis.

This approach is especially effective for events that are traditionally difficult to detect. Recent large-scale GFEs are difficult to distinguish in Ks distributions because their average Ks values are close to zero, similar to the Ks of recent paralogs and sequenced isoforms. Highlighting phylogenomically inferred paralogs facilitated easy differentiation from paralogs arising from other sources (CARY8 and CARY9 in Figure 5c). The same logic also partially safeguards against assembly artifacts such as isoforms, fragmented assemblies, and recent transcript redundancies, which are expected to contribute disproportionately to low-Ks pairwise comparisons, whereas the events emphasized here require both phylogenetic clustering of duplications at a shared node and a corresponding non-zero Ks signal across descendant taxa.

Further, we resolved other inconsistencies between phylogenetic gene-duplication mapping and Ks distributions by applying duplicate-lineage discordance (DLD) correction during gene- duplication mapping. For example, *H_2_O*’s conservative gene-duplication mapping and DLD correction resolved the confusion in Carruthers et al. (2024) about the placement of ancient WGDs along the backbone of Ericales, due to scattered gene-duplication clusters along the low- support backbone relationships. *H_2_O*’s duplication mapping is robust to alternative backbone relationships in the input consensus species tree (number change <1%). We confirmed a single large-scale GFE for Balsaminaceae (ERIC2), and a putative ancient WGD (ERIC1) for all Ericales excluding the Balsaminoids (Figure S10). Our determination aligns with the inferences from synteny analyses and assembled genomes by Zhang et al. (2020). Like ERIC1, the major large- scale GFE in Caryophyllaceae, CARY1, faces the same issue (Figure 3). It has many gene duplications clustered at two neighboring nodes, and DLD correction shifted nearly all gene duplications mapped to the parent node to the corresponding event node. This suggests that most duplicated gene families generated by these events have the homolog tree topology shown in Figure 3a, indicating they may have arisen through allopolyploidy, as in the hybridization scenario shown in the same figure. These results may reveal a new phylogenetic signature of allopolyploidy, distinct from expectations under existing reconciliation-based models of polyploidy (Thomas et al. 2017). Because the divergence times of the current lineages we view as hybrid parents are relatively short in geological terms, the phylogenetic signal of ancient hybridization may be ambiguous and could resemble ILS.

Additionally, our results suggest that Ks values show strong phylogenetic signals and may be influenced by life-history shifts and the rate of molecular evolution; thus, they are not a reliable indicator of time. Herbaceous plants experience faster rates of molecular evolution because they have shorter generation times (Smith and Donoghue 2008). Ks is a measure of molecular distance between paralogous gene pairs and is thus likely to be affected by shifts in molecular rates. Based on t-tests, the Ks peak means in herbaceous plants are statistically significantly higher compared to those in woody plants for ERIC1 (Ericales excluding Balsaminoids) and PHYT1 (Phytolaccoids) (Table 1). PHYT1 is a particularly clear example, as Nyctagineae shifted from woody to herbaceous habit, coinciding with a statistically significant increase in Ks peak means. For CARY1, Ks peak means in herbaceous and woody species do not differ significantly.

Notably, the dates for large-scale GFE events in Table 1 range from 56.7 to 110.9 Ma, but their average Ks are similar and do not correlate with age.

We inferred these dates from molecular phylogenies with fossil constraints, based on the phylogenetic placement of the large-scale GFE events, unrelated to Ks. Rather than correlating simply with time, the Ks peak means showed strong phylogenetic signals, with Pagel’s λ close to 1 for all selected events except PHYT1. This suggests that the Ks peak means contain a strong phylogenetic component consistent with Brownian-motion-like evolution. The phylogenetic ANOVA tests are mostly insignificant, potentially because the number of character shifts and/or the sample size is too small to reject the null hypothesis. The non-phylogenetic t-tests, therefore, identify patterns that may warrant further investigation, even when the available datasets have limited power for phylogenetic ANOVA. Clearly, life-history shifts and potentially other phylogenetically conserved traits may influence Ks values, indicating that Ks is not a stable measure of time. At broad taxonomic scales, life-history shifts occur frequently in angiosperm evolutionary history. Therefore, we suggest caution when making time inferences with Ks values in flowering plants.

### Mitigating Systematic Errors in Phylogenetic Orthology Inference Associated with GFEs

Large-scale GFEs/ancient WGDs are commonly associated with high levels of gene tree discordance, which is often attributed to biological causes, such as incomplete lineage sorting (ILS) and hybridization (Walker et al. 2018; Yang et al. 2018; Stull et al. 2021; Carruthers et al. 2024; Feng et al. 2024). We found that 46% of the large-scale GFEs in our three phylogenomic datasets have close descendant nodes with high discordance. These close descendant nodes of the events are referred to as GFE-associated nodes, and they have statistically significantly higher proportions of gene tree discordance than the rest of the consensus species tree across all methods (*p* < 0.0002). This suggests GFEs are a major source of gene tree discordance in the two Caryophyllales subclades and Ericales.

In addition to biological causes, our results from both simulated and empirical datasets demonstrate that systematic errors in orthology inference substantially contribute to the association between large-scale GFEs and gene tree discordance. We hypothesized that these systematic errors are caused by hidden paralogy in two ways: 1) single-copy descendant genes of gene duplications may be hidden paralogs under the influence of ILS, and 2) one-to-one homolog trees may not be true orthologs when large-scale GFEs are involved. *H_2_O* employed two phylogenetic data filtering methods to resolve these potential problems in the datasets: (1) removing single-copy descendant genes for each gene duplication (the “pruning” method; Figure 3b); and (2) removing one-to-one homologs for inferred large-scale GFE events (the “GFE” method; Figure 3c).

We simulated high levels of ILS near a hypothetical large-scale GFE event that resembles the observed level of gene tree discordance in empirical datasets. Without *H_2_O*’s two data filtering approaches, *H_2_O* unfiltered, ASTRAL-Pro3 (Zhang et al. 2025), and Yang and Smith (2014) RT and MO reported similar proportions of paralogous relationships in their inferred ortholog trees (Figure 4). This was accompanied by higher levels of gene tree discordance than those caused by the same level of ILS without gene duplication. This suggests the extra amount of discordance is driven by systematic errors in orthology inference. As intended, *H_2_O* “pruning” filtering reduced hiddenly paralogous relationships in the single-copy tips in duplicated clades, while *H_2_O* “GFE” methods removed hidden paralogs from single-copy homolog trees. However, both also removed orthologous relationships because they do not test for orthology and instead remove phylogenetic data that fit the criteria. Thus, discordance is not substantially reduced when these two methods were applied separately. However, when the two filters were applied simultaneously, *H_2_O* “pruned x GFE” reported no paralogous relationships and reduced discordance to a level even lower than that produced by ILS without gene duplication. This may be because *H_2_O* “pruned” also removed orthologous ILS relationships that conflicted with the species tree.

Similarly, in the empirical dataset, neither *H_2_O* filtering approach alone significantly changed the proportion of gene tree discordance in GFE-associated nodes, but applying both filters resulted in a statistically significant reduction (Figure 6a and 6b). Based on the simulation results, this reduction may suggest that systematic errors in orthology inference were largely removed, if not entirely. The remaining discordance in the empirical “pruned x GFE” orthologs likely reflects ILS or other methodological error rather than errors in orthology inference.

Comparing discordance across the four orthology inference methods (ASTRAL-Pro3, Yang and Smith (2014) RT and MO, *H_2_O*), single-copy homolog trees without group and homolog trees without monophyletic outgroups may not substantially contribute to gene tree discordance associated with large-scale GFEs. An important consideration regarding ASTRAL-Pro (Zhang et al. 2025) is that high posterior probability (PP) values (i.e., ∼1) do not necessarily indicate low gene tree discordance. Given the large number of input trees typical of phylogenomic datasets, PP values can remain high even when substantial gene tree discordance is present (Sayyari and Mirarab, 2016), as we also observed in the three datasets.

Filtering gene trees may recover the same or a different consensus species-tree topology. When the same topology is recovered after *H_2_O*’s substantial filtering, that agreement provides additional support for the stability of those relationships under alternative orthology treatments. Alternative topologies should be treated as hypotheses generated by a subset of the data with less systematic error but potentially still affected by ILS, e.g., ERIC1 (the backbone of Ericales, Figure S23). However, the consensus species tree topologies inferred from “pruned x GFE” ortholog trees should not be interpreted as automatically superior, because this treatment substantially reduces the amount of data available for tree inference (Figure 6c). This reduction has led to low information in relationships further from the GFE event in some groups: CARY6w (Figure S21) and ERIC1 (Figure S23). If *H_2_O*’s approaches yielded better support for relationships directly following large-scale GFE but less support for other relationships, *H_2_O* has a subcommand that can extract selected well-supported relationships around the GFE event, and these relationships can be used as a topology constraint to run ASTRAL4 with the unfiltered ortholog set to create the best-supported consensus species tree. Thus, *H_2_O* should be viewed as a conservative filtering and diagnostic approach. It can identify relationships whose inference is sensitive to hidden paralogy, but it does so at the expense of information.

Future methods that explicitly model post-GFE gene loss, hidden paralogy, and discordance without requiring such extensive data removal would be especially valuable.

### Inferred Gene Family Contractions Following GFEs Before Named Clades

Retained gene-family expansions have long been considered potential contributors to trait evolution following ancient WGD in angiosperms (Chanderbali et al. 2016; Clark and Donoghue 2018; Smith et al. 2018b; Sobhanian et al. 2026). The expansions detected in extant transcriptomic or genomic datasets are more precisely interpreted as retained duplicate-gene signals: they represent gene families for which duplicate copies remain detectable in sampled descendant taxa. Gene-family retention is known to be non-random and may reflect dosage balance, functional constraint, or adaptive advantages (Carretero-Paulet and Fares 2012; Paterson et al. 2012; Moghe et al. 2014).

To further explore inferred GFEs, phylogenomic studies often focus on gene ontology to identify which gene functions are preferentially retained, rarely examining gene families phylogenetically or investigating subsequent gene-family contractions (Carretero-Paulet and Fares 2012; Qiao et al. 2019; Wang et al. 2019b; Feng et al. 2024). In many inferred GFE- associated gene families, duplicate copies are not detected in all descendant lineages. We refer to these lineage-specific reductions in detectable copy number as “gene-family contractions”.

In transcriptomic datasets, the absence of a transcript may reflect true genomic loss, lack of expression in sampled tissues, sequencing/assembly quality, or filtering during homolog construction. These inferred reductions in duplicate-gene detectability have received less attention than duplicate retention in studies of angiosperm genome evolution. In existing studies, the focus is often at the genome level, such as the physiological and ecological benefits of genome downsizing (Dodsworth et al. 2016; Simonin and Roddy 2018; Wang et al. 2021), or on probabilistic modeling of gene copy loss (Tiley et al. 2016; Ren et al. 2018; Zwaenepoel and Van de Peer 2019). Here, we use the phylogenetic distribution of undetected duplicate copies to explore whether post-GFE reductions in detectable copy number show non-random patterns across clades.

In the three phylotranscriptomic datasets, molecular branch length is weakly to moderately correlated with the size of contractions (Figure S26). Molecular branch length reflects the rate of molecular evolution and time. Thus, longer branch lengths may indicate more time for gene family contractions to occur and may correlate positively with contraction size. The varying correlations observed could reflect heterogeneity in molecular rates, data quality, and branch- length inference issues. A relevant issue in branch-length inference is spurious substitutions produced by ILS (SPILS; Mendes and Hahn 2016). This phenomenon occurs because some relationships in discordant ILS gene trees are absent in the consensus species tree. The substitutions on those branches need to be placed elsewhere, leading to spuriously longer or shorter branch lengths near the discordant relationship. Therefore, when large-scale GFEs are associated with strong discordance in nearby nested nodes, neighboring branches may appear falsely longer due to SPILS, creating an artificial correlation between gene-family contraction and molecular branch length.

We also observed that larger inferred contractions often map to branches immediately preceding named clades, i.e., families, tribes, and genera (Figure 7a). Although many large-scale GFEs have been associated with major clades and trait innovations (Chanderbali et al. 2016; Clark and Donoghue 2018; Smith et al. 2018b; Yang et al. 2018; Sobhanian et al. 2026), our results suggest that reductions in detectable duplicate-gene content may also be concentrated on branches preceding taxonomically recognized clades. This pattern suggests that post-GFE duplicate loss, expression loss, or reduced transcript detectability may accompany some of the morphological or ecological shifts used to recognize these groups. For instance, among the gene families inferred to have been retained after the putative ancient WGD ERIC1, 29% are not detected along the branch leading to the MRCA of the parasitic genera *Eremotropa* and *Monotropa*. This pattern is consistent with, but does not by itself demonstrate, gene loss associated with the reduction or loss of photosynthetic function in these lineages (Gruzdev et al. 2016; Ravin et al. 2016). These patterns suggest that post-GFE reductions in duplicate-gene content deserve greater attention alongside duplicate retention, particularly in clades with major shifts in ecology, morphology, or life history. However, after correcting for molecular branch length, the relationship between gene family contractions and named clades is no longer statistically significant. This correction may not be justified if the correlation between branch lengths and the size of gene family contractions is spurious, e.g., due to SPILS. Testing whether these inferred contractions reflect true genomic losses, expression differences, or transcriptomic sampling effects will require genome-scale data, broader tissue sampling, and explicit models of duplicate retention and loss. Further investigation is also needed to assess the extent of SPILS and the possible influence of inferred duplicate-copy loss on trait evolution.

### Conclusions and Future Directions

Large-scale gene family expansions (GFEs), including WGDs, chromosomal duplications, and segmental duplications, have played important roles in flowering plant evolution, but detecting and placing them remains challenging when analyses rely primarily on transcriptomic data. Our results show that node-partitioned Ks distributions improve confidence in the placement of inferred GFEs and clarify the limits of transcriptome-based inference. Accurate detection of these evolutionarily important events strengthens the analytical framework underlying downstream studies of adaptation and molecular pathways associated with large-scale GFEs.

Our command-line tool, *H_2_O*, provides a framework for jointly evaluating Ks evidence, mapped gene duplications, and orthology inference in clades affected by ancient duplication. Applied to simulated data and three empirical datasets, this framework identified GFE-associated regions with elevated gene-tree discordance and showed that filtering gene trees likely affected by hidden paralogy can significantly reduce discordance induced by errors in orthology inference. Because this reduction is accompanied by substantial data loss, consensus species trees inferred from filtered topologies should be interpreted as conservative hypotheses rather than superior. Nevertheless, these findings emphasize the importance of accounting for hidden paralogy in phylogenetic orthology inference, particularly in clades with histories of large-scale GFEs and hybridization. Given the widespread occurrence of both processes across angiosperms, *H_2_O* provides a broadly applicable framework for improving phylogenomic inference in flowering plants and is generalizable to datasets combining transcriptomes and genomes.

Our results also suggest that ancient hybridization may, in some cases, generate gene-tree patterns that are difficult to distinguish from ILS. In addition, Ks values vary with life history and exhibit strong phylogenetic signal, underscoring the need for caution when using Ks as a proxy for time across lineages with different molecular rates. Recent studies have increasingly used Ks values to date large-scale GFE events and infer broader evolutionary hypotheses (Chen et al. 2024; Chen et al. 2026). Nevertheless, Ks is approximately proportional to the product of divergence time and lineage-specific substitution rates; however, no reliable method exists to extract substitution rates from the number of substitutions in Ks (Smith and Beaulieu 2024). Consequently, temporal interpretations based on Ks should be made cautiously, particularly when comparing lineages with different molecular evolutionary rates. Finally, patterns of inferred duplicate-copy absence following GFEs suggest that post-GFE reductions in detectable gene-family content may be an important, but still underexplored, component of angiosperm genome evolution.

More broadly, these results emphasize that marker choice matters for studies of genome evolution. Datasets enriched for apparent single-copy loci remain valuable for many phylogenetic questions, but they may be less informative for detecting the history and consequences of gene-family expansion. Expanded genome and transcriptome sampling, combined with methods that explicitly account for duplication, loss, expression, and hidden paralogy, will be essential for reconstructing the genomic history of flowering plants.

## Data Availability

The package *H_2_O* is available at https://github.com/fkyeeb/H2O. Processed homolog trees, ortholog trees, scripts, and other data files are available at https://doi.org/10.5281/zenodo.20347899.

## Author Contributions

K.F., M.L.G., and S.A.S. designed the research; K.F. analyzed the data and led the writing; M.L.G. and S.A.S. assisted with data interpretation and the writing.

## Acknowledgments

K.F. and S.A.S. were supported in part by National Science Foundation grants, including IntBio 2217116, DEB-NERC 1939226, DBI 1930030, and M.L.G. by the Postdoctoral Research Fellowship in Biology (DBI-2410238). We thank Joseph Walker for discussions on the initial ideas; Rebekah Mohn for discussions on ancient hybridization; and James Boyko, Joey Charboneau, Brad Ruhfel, Will Weaver, and Avinash Subramanian for helpful discussions on the manuscript. We also thank the editors of MBE and three anonymous reviewers for constructive comments that substantially improved the manuscript.

## Reference

1. Amborella Genome Project, Albert VA, Barbazuk WB, de Pamphilis CW, Der JP, Leebens-Mack J, Ma H, Palmer JD, Rounsley S, Sankoff D, et al. 2013. The *Amborella* genome and the evolution of flowering plants. Science 342:1241089.

2. Ballesteros JA, Hormiga G. 2016. A new orthology assessment method for phylogenomic data: Unrooted phylogenetic orthology. Mol Biol Evol. 33:2117–2134.

3. Brown JW, Walker JF, Smith SA. 2017. Phyx: Phylogenetic tools for unix. Bioinformatics 33:1886–1888.

4. Cai L, Xi Z, Amorim AM, Sugumaran M, Rest JS, Liu L, Davis CC. 2019. Widespread ancient whole-genome duplications in Malpighiales coincide with Eocene global climatic upheaval. New Phytol. 221:565–576.

5. Carretero-Paulet L, Fares MA. 2012. Evolutionary dynamics and functional specialization of plant paralogs formed by whole and small-scale genome duplications. Mol Biol Evol. 29:3541–3551.

6. Carruthers T, Brockington SF, Maeda HA, Stull GW, Walker-Hale N, Yang Y, Smith SA. 2026. Evolution in arid environments: the role of betalains, succulence, and medullary bundles. New Phytol. 252:1250–1268.

7. Carruthers T, Gonçalves DJP, Li P, Chanderbali AS, Dick CW, Fritsch PW, Larson DA, Soltis DE, Soltis PS, Weaver WN, et al. 2024. Repeated shifts out of tropical climates preceded by whole genome duplication. New Phytol. 244:2561–2575.

8. Chanderbali AS, Berger BA, Howarth DG, Soltis PS, Soltis DE. 2016. Evolving ideas on the origin and evolution of flowers: New perspectives in the genomic era. Genetics 202:1255–1265.

9. Chen H, Almeida-Silva F, Logghe G, Maere S, Bonte D, Van de Peer Y. 2026. The rise of polyploids during environmental upheaval. Cell 189:3608–3622.e5.

10. Chen D, Zhang T, Chen Y, Ma H, Qi J. 2022. Tree2GD: a phylogenomic method to detect large- scale gene duplication events. Bioinformatics 38:5317–5321.

11. Chen H, Zwaenepoel A, Van de Peer Y. 2024. wgd v2: a suite of tools to uncover and date ancient polyploidy and whole-genome duplication. Bioinformatics 40:btae272.

12. Clark JW, Donoghue PCJ. 2018. Whole-genome duplication and plant macroevolution. Trends Plant Sci. 23:933–945.

13. Degnan JH, Rosenberg NA. 2009. Gene tree discordance, phylogenetic inference and the multispecies coalescent. Trends in Ecology & Evolution 24:332–340.

14. Dodsworth S, Chase MW, Leitch AR. 2016. Is post-polyploidization diploidization the key to the evolutionary success of angiosperms? Bot J Linn Soc 180:1–5.

15. Feng K, Walker JF, Marx HE, Yang Y, Brockington SF, Moore MJ, Rabeler RK, Smith SA. 2024. The link between ancient whole-genome duplications and cold adaptations in the Caryophyllaceae. Amer J Bot. 111:e16350.

16. Feng T, McKibben M, Lovell J, Michelmore R, Rieseberg LH, Barker MS, Schranz ME. 2026. Phylogenomic synteny reveals paleohexaploid-derived genomic blocks across Asteraceae. PNAS 123:e2426851123.

17. Fitch WM. 1970. Distinguishing Homologous from Analogous Proteins. Syst Biol. 19:99–113.

18. Galtier N, Daubin V. 2008. Dealing with incongruence in phylogenomic analyses. Philos Trans R Soc B: Biol Sci. 363:4023–4029.

19. García N, Folk RA, Meerow AW, Chamala S, Gitzendanner MA, Oliveira RS de, Soltis DE, Soltis PS. 2017. Deep reticulation and incomplete lineage sorting obscure the diploid phylogeny of rain-lilies and allies (Amaryllidaceae tribe Hippeastreae). Mol Phylogenet Evol. 111:231–247.

20. Gaynor ML, Feng K, Soltis DE, Soltis PS, Smith SA, unpublished data, 10.64898/2026.08.25.747000, last accessed November 16, 2026.

21. Gruzdev EV, Mardanov AV, Beletsky AV, Kochieva EZ, Ravin NV, Skryabin KG. 2016. The complete chloroplast genome of parasitic flowering plant *Monotropa hypopitys*: Extensive gene losses and size reduction. Mitochondrial DNA B Resour. 1:212–213.

22. Heslop-Harrison JS (Pat), Schwarzacher T, Liu Q. 2023. Polyploidy: Its consequences and enabling role in plant diversification and evolution. Ann Bot. 131:1–10.

23. Huang C-H, Zhang C, Liu M, Hu Y, Gao T, Qi J, Ma H. 2016. Multiple polyploidization events across Asteraceae with two nested events in the early history revealed by nuclear phylogenomics. Mol Biol Evol. 33:2820–2835.

24. Jiao Y, Li J, Tang H, Paterson AH. 2014. Integrated syntenic and phylogenomic analyses reveal an ancient genome duplication in Monocots. Plant Cell 26:2792–2802.

25. Kozlov AM, Darriba D, Flouri T, Morel B, Stamatakis A. 2019. RAxML-NG: a fast, scalable and user-friendly tool for maximum likelihood phylogenetic inference. Bioinformatics 35:4453– 4455.

26. Landis JB, Soltis DE, Li Z, Marx HE, Barker MS, Tank DC, Soltis PS. 2018. Impact of whole-genome duplication events on diversification rates in angiosperms. Amer J Bot. 105:348–363.

27. Leebens-Mack JH, Barker MS, Carpenter EJ, Deyholos MK, Gitzendanner MA, Graham SW, Grosse I, Li Z, Melkonian M, Mirarab S, et al. 2019. One thousand plant transcriptomes and the phylogenomics of green plants. Nature 574:679–685.

28. Li Z, McKibben MTW, Finch GS, Blischak PD, Sutherland BL, Barker MS. 2021. Patterns and processes of diploidization in land plants. Annu Rev Plant Biol. 72:387–410.

29. Liu Z, Li M, Wang Y, Li T, Li W, Xing L, Du H, He Q. 2025. Genomic insights into karyotype evolution and adaptive mechanisms in Polygonaceae species. Cell Reports 44.

30. Mallo D, De Oliveira Martins L, Posada D. 2016. SimPhy: Phylogenomic Simulation of Gene, Locus, and Species Trees. Syst Biol 65:334–344.

31. Marcet-Houben M, Gabaldón T. 2011. TreeKO: A duplication-aware algorithm for the comparison of phylogenetic trees. Nucleic Acids Res 39:e66.

32. McKain MR, Tang H, McNeal JR, Ayyampalayam S, Davis JI, de Pamphilis CW, Givnish TJ, Pires JC, Stevenson DWm, Leebens-Mack JH. 2016. A phylogenomic assessment of ancient polyploidy and genome evolution across the Poales. Genome Biol Evol 8:1150–1164.

33. Mendes FK, Hahn MW. 2016. Gene tree discordance causes apparent substitution rate variation. Syst Biol. 65:711–721.

34. Moghe GD, Hufnagel DE, Tang H, Xiao Y, Dworkin I, Town CD, Conner JK, Shiu S-H. 2014. Consequences of whole-genome triplication as revealed by comparative genomic analyses of the Wild Radish *Raphanus raphanistrum* and three other Brassicaceae species. Plant Cell 26:1925–1937.

35. Paterson AH, Wang X, Li J, Tang H. 2012. Ancient and recent polyploidy in Monocots. In: Soltis PS, Soltis DE, editors. Polyploidy and genome evolution. Berlin, Heidelberg: Springer. p. 93–108.

36. POWO. 2026. Plants of the world online. Facilitated by the Royal Botanic Gardens, Kew. Published on the Internet. Website: http://www.plantsoftheworldonline.org/. [accessed January 2026].

37. Qiao X, Li Q, Yin H, Qi K, Li L, Wang R, Zhang S, Paterson AH. 2019. Gene duplication and evolution in recurring polyploidization–diploidization cycles in plants. Genome Biol 20:38.

38. Ravin NV, Gruzdev EV, Beletsky AV, Mazur AM, Prokhortchouk EB, Filyushin MA, Kochieva EZ, Kadnikov VV, Mardanov AV, Skryabin KG. 2016. The loss of photosynthetic pathways in the plastid and nuclear genomes of the non-photosynthetic mycoheterotrophic eudicot *Monotropa hypopitys*. BMC Plant Biol 16:238.

39. Ren R, Wang H, Guo C, Zhang N, Zeng L, Chen Y, Ma H, Qi J. 2018. Widespread whole genome duplications contribute to genome complexity and species diversity in Angiosperms. Mol Plant. 11:414–428.

40. Revell LJ. 2024. phytools 2.0: An updated R ecosystem for phylogenetic comparative methods (and other things). PeerJ 12:e16505.

41. Rice A, Glick L, Abadi S, Einhorn M, Kopelman NM, Salman-Minkov A, Mayzel J, Chay O, Mayrose I. 2015. The Chromosome Counts Database (CCDB) – A community resource of plant chromosome numbers. New Phytol. 206:19–26.

42. Salmon A, Hao Y, Milin M, Lima O, Cavé-Radet A, Giraud D, Cruaud C, Labadie K, Istace B, Belser C, et al. 2025. On the way to diploidization and unexpected ploidy in the grass *Sporobolus* section *Spartina* mesopolyploids. Nat Commun 16:1997.

43. Sayyari E, Mirarab S. 2016. Fast coalescent-based computation of local branch support from quartet frequencies. Mol Biol Evol. 33:1654–1668.

44. Simonin KA, Roddy AB. 2018. Genome downsizing, physiological novelty, and the global dominance of flowering plants. PLOS Biology 16:e2003706.

45. Smith SA, Beaulieu JM. 2024. *Ad fontes*: Divergence-time estimation and the age of angiosperms. New Phytol. 244:760–766.

46. Smith SA, Brown JW, Walker JF. 2018. So many genes, so little time: A practical approach to divergence-time estimation in the genomic era. PLoS One 13:e0197433.

47. Smith SA, Brown JW, Yang Y, Bruenn R, Drummond CP, Brockington SF, Walker JF, Last N, Douglas NA, Moore MJ. 2018. Disparity, diversity, and duplications in the Caryophyllales. New Phytol. 217:836–854.

48. Smith SA, Donoghue MJ. 2008. Rates of molecular evolution are linked to life history in flowering plants. Science 322:86–89.

49. Smith SA, Moore MJ, Brown JW, Yang Y. 2015. Analysis of phylogenomic datasets reveals conflict, concordance, and gene duplications with examples from animals and plants. BMC Evolutionary Biology 15:150.

50. Smith SA, Pease JB, Carruthers T, Bradburd GS, Huegele IB, Stull GW, Weaver WN, Yang Y, Yi T- S, Beaulieu JM. 2026. Longevity in plants impacts phylogenetic and population dynamics. New Phytol. 250:661–671.

51. Smith SA, Walker-Hale N, Walker JF, Brown JW. 2020. Phylogenetic conflicts, combinability, and deep phylogenomics in plants. Syst Biol. 69:579–592.

52. Sobhanian H, Song W-Y, Soltis PS, Soltis DE, Chen S. 2026. Polyploidy and plant resilience to environmental stresses: Molecular mechanisms and future applications. Plant Comm 7.

53. Soltis DE, Visger CJ, Soltis PS. 2014. The polyploidy revolution then…and now: Stebbins revisited. Amer J Bot. 101:1057–1078.

54. Stull GW, Pham KK, Soltis PS, Soltis DE. 2023. Deep reticulation: The long legacy of hybridization in vascular plant evolution. Plant J. 114:743–766.

55. Stull GW, Qu X-J, Parins-Fukuchi C, Yang Y-Y, Yang J-B, Yang Z-Y, Hu Y, Ma H, Soltis PS, Soltis DE, et al. 2021. Gene duplications and phylogenomic conflict underlie major pulses of phenotypic evolution in gymnosperms. Nat Plants. 7:1015–1025.

56. Tabatabaee Y, Zhang C, Warnow T, Mirarab S. 2023. Phylogenomic branch length estimation using quartets. Bioinformatics 39:i185–i193.

57. Tang H, Wang X, Bowers JE, Ming R, Alam M, Paterson AH. 2008. Unraveling ancient hexaploidy through multiply-aligned angiosperm gene maps. Genome Res. 18:1944–1954.

58. Thomas GWC, Ather SH, Hahn MW. 2017. Gene-tree reconciliation with MUL-trees to resolve polyploidy events. Syst Biol. 66:1007–1018.

59. Tiley GP, Ané C, Burleigh JG. 2016. Evaluating and characterizing ancient whole-genome duplications in plants with gene count data. Genome Biol Evol 8:1023–1037.

60. Tiley GP, Barker MS, Burleigh JG. 2018. Assessing the performance of Ks plots for detecting ancient whole genome duplications. Genome Biol Evol 10:2882–2898.

61. Triplett JK, Clark LG, Fisher AE, Wen J. 2014. Independent allopolyploidization events preceded speciation in the temperate and tropical woody bamboos. New Phytol. 204:66–73.

62. Walker JF, Yang Y, Feng T, Timoneda A, Mikenas J, Hutchison V, Edwards C, Wang N, Ahluwalia S, Olivieri J, et al. 2018. From cacti to carnivores: Improved phylotranscriptomic sampling and hierarchical homology inference provide further insight into the evolution of Caryophyllales. Amer J Bot. 105:446–462.

63. Walker JF, Yang Y, Moore MJ, Mikenas J, Timoneda A, Brockington SF, Smith SA. 2017. Widespread paleopolyploidy, gene tree conflict, and recalcitrant relationships among the carnivorous Caryophyllales. Amer J Bot. 104:858–867.

64. Wang J, Qin J, Sun P, Ma X, Yu J, Li Y, Sun S, Lei T, Meng F, Wei C, et al. 2019. Polyploidy Index and its implications for the evolution of polyploids. Front Genet. 10.

65. Wang N, Yang Y, Moore MJ, Brockington SF, Walker JF, Brown JW, Liang B, Feng T, Edwards C, Mikenas J, et al. 2019. Evolution of Portulacineae marked by gene tree conflict and gene family expansion associated with adaptation to harsh environments. Mol Biol Evol. 36:112–126.

66. Wang X, Morton JA, Pellicer J, Leitch IJ, Leitch AR. 2021. Genome downsizing after polyploidy: Mechanisms, rates and selection pressures. Plant J. 107:1003–1015.

67. Yang Y, Moore MJ, Brockington SF, Mikenas J, Olivieri J, Walker JF, Smith SA. 2018. Improved transcriptome sampling pinpoints 26 ancient and more recent polyploidy events in Caryophyllales, including two allopolyploidy events. New Phytol. 217:855–870.

68. Yang Y, Moore MJ, Brockington SF, Soltis DE, Wong GK-S, Carpenter EJ, Zhang Y, Chen L, Yan Z, Xie Y, et al. 2015. Dissecting molecular evolution in the highly diverse plant clade Caryophyllales using transcriptome sequencing. Mol Biol Evol. 32:2001–2014.

69. Yang Y, Smith SA. 2014. Orthology inference in nonmodel organisms using transcriptomes and low-coverage genomes: Improving accuracy and matrix occupancy for phylogenomics. Mol Biol Evol. 31:3081–3092.

70. Zhang C, Nielsen R, Mirarab S. 2025. ASTER: A package for large-scale phylogenomic reconstructions. Mol Biol Evol. 42:msaf172.

71. Zhang C, Scornavacca C, Molloy EK, Mirarab S. 2020. ASTRAL-Pro: Quartet-based species-tree inference despite paralogy. Mol Biol Evol. 37:3292–3307.

72. Zhang C, Zhang T, Luebert F, Xiang Y, Huang C-H, Hu Y, Rees M, Frohlich MW, Qi J, Weigend M, et al. 2020. Asterid phylogenomics/phylotranscriptomics uncover morphological evolutionary histories and support phylogenetic placement for numerous whole-genome duplications. Mol Biol Evol. 37:3188–3210.

73. Zhao L, Yang Y-Y, Qu X-J, Ma H, Hu Y, Li H-T, Yi T-S, Li D-Z. 2023. Phylotranscriptomic analyses reveal multiple whole-genome duplication events, the history of diversification and adaptations in the Araceae. Ann Bot. 131:199–214.

74. Zwaenepoel A, Van de Peer Y. 2019. Inference of ancient whole-genome duplications and the evolution of gene duplication and loss rates. Mol Biol Evol. 36:1384–1404.

